# Negative feedback of cyclic di-GMP levels optimizes switching between sessile and motile lifestyles in *Vibrio cholerae*

**DOI:** 10.1101/2024.09.01.610008

**Authors:** Aathmaja Anandhi Rangarajan, Jeremy W. Schroeder, Rebecca L. Hurto, Geoffrey B. Severin, Macy E. Pell, Meng-Lun Hsieh, Christopher M. Waters, Lydia Freddolino

**Affiliations:** Department of Microbiology, Genetics and Immunology, Michigan State University, East Lansing, MI, USA; Department of Biological Chemistry, University of Michigan, Ann Arbor, MI 48109, USA; Department of Computational Medicine and Bioinformatics, University of Michigan, Ann Arbor, MI 48109, USA

## Abstract

The signaling molecule cyclic di-GMP (c-di-GMP) controls the switch between bacterial motility and biofilm production, and fluctuations in cellular levels of c-di-GMP have been implicated in *Vibrio cholerae* pathogenesis. Intracellular concentrations of c-di-GMP are controlled by the interplay of diguanylate cyclase (DGC) enzymes, which synthesize c-di-GMP to promote biofilms, and phosphodiesterase (PDE) enzymes, which hydrolyze c-di-GMP to drive motility. To track the complete regulatory logic of how *V. cholerae* responds to changing c-di-GMP levels, we followed a time course of overexpression of either the *V. campbellii* diguanylate cyclase QrgB or a variant of QrgB lacking catalytic activity (QrgB*). We find that QrgB increases c-di-GMP levels relative to QrgB* for 30 minutes after overexpression, but the effect of QrgB on c-di-GMP levels plateaus at 30 minutes, indicating tight adaptive control of c-di-GMP levels. In contrast, loss of VpsR, a master regulator activating biofilm formation upon binding to c-di-GMP, leads to higher baseline levels of c-di-GMP and continuously increasing c-di-GMP through 60 minutes after QrgB induction, revealing the existence of a negative feedback loop on c-di-GMP levels operating through VpsR. Through a combination of RNA polymerase ChIP-seq, RNA-seq, and genetic approaches, we show that transcription of a gene encoding a PDE, *cdgC*, is activated by VpsR at high c-di-GMP concentrations, mediating this negative feedback on c-di-GMP levels. Further, although cells lacking *cdgC* exhibit enhanced biofilm formation, these mutants are outcompeted by wild type *V. cholerae* in colonization assays that reward a combination of attachment, dispersal, and motility behaviors. These results underscore the importance of negative feedback regulation of c-di-GMP to maintain appropriate homeostatic levels for efficient transitioning between biofilm formation and motility, both of which are necessary over the course of the *V. cholerae* infection cycle.

## Introduction

*Vibrio cholerae* is the causative agent of cholera, a severe diarrheal disease which affects millions of people and causes nearly 100,000 deaths annually, with the number of deaths increasing significantly in recent years according to WHO reports^1^. *V. cholerae* can occupy (and transition between) numerous environmental niches including metazoan digestive tracts, arthropod exoskeletons, and aquatic habitats, requiring rapid lifestyle shifts between a sessile, biofilm state and a motile, free-living state for successful survival and propagation^2–5^. In the majority of bacteria, an important signaling molecule that directs the transition between motility and biofilms is the second messenger cyclic di-GMP (c-di-GMP), with high c-di-GMP universally driving bacteria toward biofilm association and low c-di-GMP promoting unicellular lifestyles^3,5,6^.

c-di-GMP is recognized by numerous transcription factors, riboswitches, and sRNAs to regulate gene expression both transcriptionally and post-transcriptionally^6,7^. In *V. cholerae*, two master regulators promote biofilm formation when they bind c-di-GMP: VpsR and VpsT^8,9^. c-di-GMP-bound VpsR activates transcription of *vpsT*, and both VpsR and VpsT activate the production of biofilm matrix components when they are bound to c-di-GMP, including *vpsL*, *vpsU, rbmAC, rbmF,* and *bap-1*^10–17^. At the same time, c-di-GMP binds to and inactivates the master regulator of flagellar gene expression, FlrA^18,19^, to reduce flagellar motility.

Several mechanistic aspects of the switch from biofilm formation to dispersal in *V. cholerae* have been identified. For example, low levels of c-di-GMP enhance the retraction of MSHA pili, detachment from surfaces through the LapGD module, downregulation of extracellular polysaccharide synthesis genes, and activation of FlrA to promote expression of flagellar genes, altogether resulting in detachment from surfaces, dispersal from biofilms, and increased motility^3,4,20,21^. However, the upstream regulatory logic affecting c-di-GMP metabolism and directing these molecular events during the switch from biofilm formation to dispersal is poorly understood.

To define the regulatory paths connecting c-di-GMP levels to the multitude of phenotypes described above, we applied a series of genome-wide studies on gene regulation and expression upon strong induction of a non-native diguanylate cyclase (DGC), *Vibrio campbellii* QrgB. We identified a genetic circuit that is important for negative feedback on c-di-GMP levels to balance biofilm formation and dispersal. Our results highlight an unexpectedly central role of regulation of the phosphodiesterase (PDE) CdgC, permitting precise adaptation of cellular c-di-GMP to maximize responsiveness to new stimuli. The regulation of c-di-GMP levels by *cdgC* highlights the importance of negative feedback regulation in *V. cholerae* to maintain c-di-GMP homeostasis at an appropriate level for optimal transitioning between biofilm formation and dispersal.

## Results

### Feedback regulation of c-di-GMP levels drives time-dependent regulation of flagellar and biofilm genes

To map the c-di-GMP dependent transcriptome of *V. cholerae* and determine the dynamics of gene regulation in response to increased c-di-GMP production, we performed a time course of induction of c-di-GMP synthesis using QrgB, a well characterized DGC from *V. campbellii* which is known to induce physiologically relevant c-di-GMP levels in *V. cholerae*^22,23^. QrgB is a cytoplasmic protein containing an N-terminal sensor PAS domain and C-terminal GGEEF domain. It also contains an RxxD I-site in the GGEEF domain that binds c-di-GMP. QrgB does not have any detectable homology to *V. cholerae* proteins, which reduces the possibility of interaction with *V. cholerae* proteins. Unless otherwise noted, all experiments were performed in a Δ*vpsL* background to avoid clumping of cells due to biofilm formation; thus, all strains below (including the “wild type”) are Δ*vpsL* mutants unless otherwise indicated. Samples were harvested at 0, 15, 30, and 60 minutes after induction of QrgB. At each timepoint, we also harvested equivalent samples from control cells expressing QrgB*, which is a catalytically inactive variant of QrgB which cannot produce c-di-GMP. At each timepoint we measured intracellular levels of c-di-GMP and performed RNA-seq and RNA polymerase (RNAP) ChIP-seq. Identical experiments were performed using Δ*vpsR* and Δ*vpsT* mutants to determine the impact of these transcription factors on the c-di-GMP dependent transcriptome.

In wild type and Δ*vpsT*, c-di-GMP levels increased at 15 and 30 minutes post-induction of QrgB relative to QrgB*, but plateaued at 60 minutes (**Fig. 1A-C**), indicating that even upon overexpression of a DGC, *V. cholerae* has mechanisms to limit c-di-GMP accumulation. In contrast, Δ*vpsR* cells had a higher concentration of c-di-GMP at the initial timepoint, and this concentration remained elevated at all timepoints when compared to wild type and Δ*vpsT* cells (**Fig. 1A-B**). Induction of the inactive QrgB* variant did not increase c-di-GMP levels in any background (**Fig. 1C**); rather, c-di-GMP concentrations steadily decreased over the course of growth in QrgB*-expression cells. To address whether this reduction in c-di-GMP was due to expression of QrgB*, we transformed a vector control plasmid into the WT strain and observed that the c-di-GMP concentrations after the addition of IPTG decreased over time similar to QrgB* **(Fig. S1A).** Similarly, there is no significant difference in cell volume of any strains at any time points (**Fig. S1B**). These results indicate that expression of QrgB* does not decrease the c-di-GMP levels or availability, but rather this decrease is due to the cells transitioning into the high cell density quorum state, consistent with the known reduction of c-di-GMP levels at high cell density^22,24^. These results suggest that c-di-GMP levels are regulated by a negative feedback loop that depends on VpsR.

**Figure 1.**
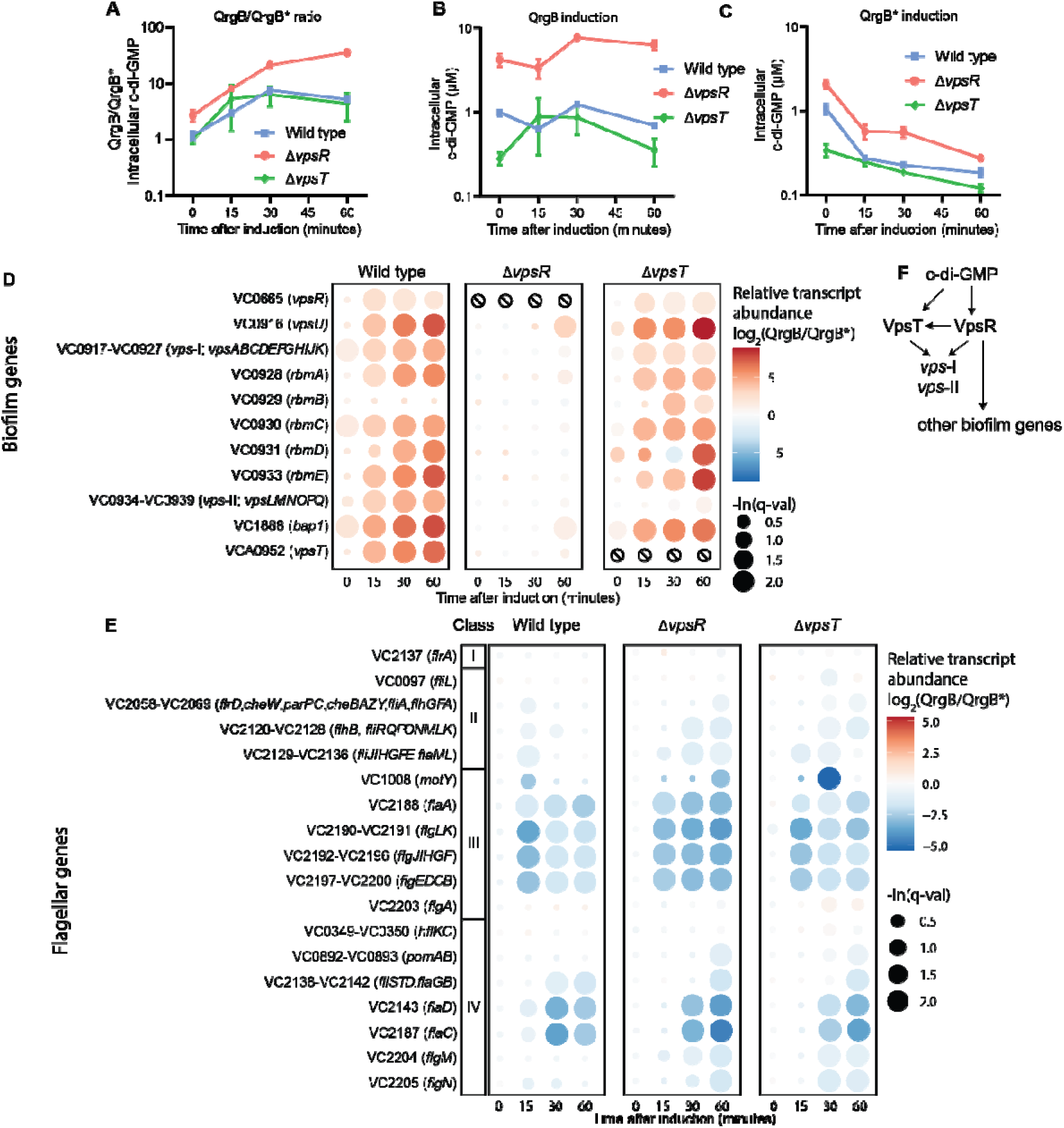
VpsR is involved in negative feedback regulation of c-di-GMP levels. In all cases QrgB and QrgB* were induced with 1mM IPTG at an OD of 1 beginning at 0 min. **A)** Data in panels A and B are represented as a ratio of c-di-GMP levels in QrgB/QrgB* over time. **B,C)** Intracellular concentrations of c-di-GMP measured by UPLC-MS/MS in WT, Δ*vpsR*, and Δ*vpsT* mutants at different timepoints after QrgB **(B)** or QrgB* **(C)** induction. Points and error bars in panels A-C represent the mean ± SEM of 3 biological replicates taking the ratio of c-di-GMP levels upon induction of QrgB vs. QrgB*. **D)** RNA-seq results for biofilm-related genes and operons. Color scale denotes the log_2_-scaled ratio between expression in QrgB induction versus QrgB* induction, and the size of each point indicates the false discovery rate, with the largest points denoting a q-value <= 0.1 and the smallest points denoting a q-value = 1.0. Null symbols () indicate that data are not available for the row in question since that gene was deleted. **E)** RNA-seq results for flagella-related genes and operons. Color scale and size of points are the same as panel C, except the color scale limits differ between panels C and D. “Class” indicates the step in the flagellum synthesis cascade in which the indicated operon has been assigned by ^45^. **F)** Regulatory network implied by our RNA-seq results.

We next examined the temporal dynamics of gene expression in response to c-di-GMP. At each timepoint we compared transcript abundance (measured via RNA-seq) following induction of QrgB vs. QrgB*. In broad terms, this experiment captured the known biological responses to c-di-GMP, as exemplified by both a gradual increase in the expression of biofilm genes over the induction (**Fig. 1D**) and a reduction in expression of flagellar genes immediately following induction (**Fig. 1E**).

Closer inspection of trends in c-di-GMP transcriptome dynamics revealed biofilm gene induction to be entirely dependent on VpsR, as expected, whereas VpsT was required for a subset of transcriptional changes including induction of the *vps*-II operon and enhanced induction of the *vps-*I operon (**Fig. 1D, F**). Flagellar expression dynamics provided further evidence of a negative feedback loop on c-di-GMP levels that depend on VpsR. Flagellar genes were repressed in a pulsatile manner in wild type cells, with c-di-GMP leading to the strongest repression of class III flagellar genes at 15 minutes (**Fig. 1E**), and class IV genes at 30 minutes. By contrast, the Δ*vpsR* mutant exhibited continuously strengthening repression of class III and IV flagellar genes throughout our time course (**Fig. 1E**), closely tracking the changes in c-di-GMP levels shown in **Fig. 1C**. The Δ*vpsT* mutant exhibited complex dynamics of flagellar gene expression resembling the wild type strain for the class III genes and the Δ*vpsR* mutant for the class IV genes. Together, these data suggest that a VpsR-dependent negative feedback loop centered on c-di-GMP acts to limit both c-di-GMP levels and flagella gene repression during the c-di-GMP response.

### CdgC is regulated by VpsR in the presence of c-di-GMP

The *V. cholerae* genome strain C6706 contains 62 enzymes predicted to encode DGCs or PDEs^25^. This includes 31 proteins with GGDEF domains involved in synthesis of c-di-GMP; 12 and 9 with EAL and HD-GYP domains, respectively, each of which could hydrolyze c-di-GMP; and 10 proteins containing both GGDEF and EAL domains [reviewed in ^6^]. To identify the gene responsible for the VpsR-dependent negative feedback on c-di-GMP levels, we assessed the effect of c-di-GMP induction on expression of transcripts encoding proteins with GGDEF, EAL, or HD-GYP domains in our RNA-seq datasets.

The GGDEF/EAL domain containing protein CdgC stood out as the most obvious candidate for negative feedback on c-di-GMP levels as it uniquely displayed c-di-GMP and time-dependent upregulation in wild type and Δ*vpsT* cells when comparing QrgB to QrgB* that was absent in the Δ*vpsR* mutant (**Fig. 2**). Although there is an apparent positive spike in the effect of c-di-GMP on CdgC expression after 15 minutes of induction in Δ*vpsR* cells, we note that this apparent increase does not represent increased expression in the QrgB condition per se, but rather, in the QrgB* condition, Δ*vpsR* cells display very low CdgC expression at this timepoint (**Fig. S2A and S2B;** see also RT-qPCR data below). Our results are consistent with previous studies that have demonstrated CdgC expression to be downregulated in a *vpsR* mutant compared to wild type, and the *cdgC* promoter to possess a VpsR binding motif [11,13]. In addition, RNAP ChIP-seq showed that increasing c-di-GMP levels led to clear increases in RNAP occupancy immediately 5′ to *cdgC* specifically in wild type cells but not in Δ*vpsR* cells (**Fig. 3A**).

**Figure 2.**
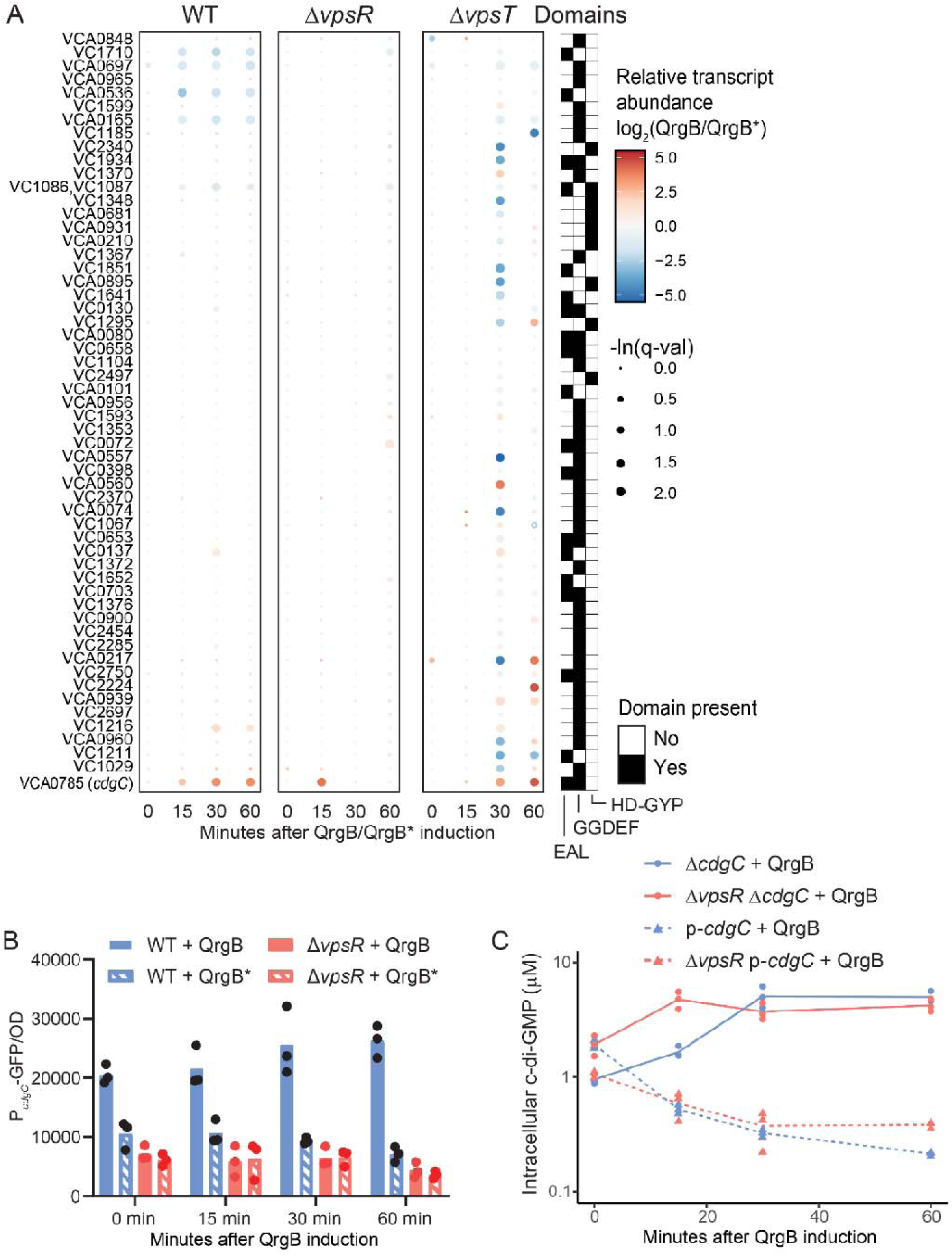
CdgC is responsible for reduction in c-di-GMP levels during c-di-GMP response. **A)** Effects of c-di-GMP induction on transcript abundance for all genes with EAL, GGDEF, or HD-GYP domains. Point size indicates statistical significance, with larger points denoting lower FDR (with the largest points for FDR < 0.1), and the color scale denotes the log_2_-scaled fold difference between gene expression in QrgB vs. QrgB*. Heatmap rows were ordered by increasing effect of c-di-GMP in WT cells at 60 minutes.

**Figure 3:**
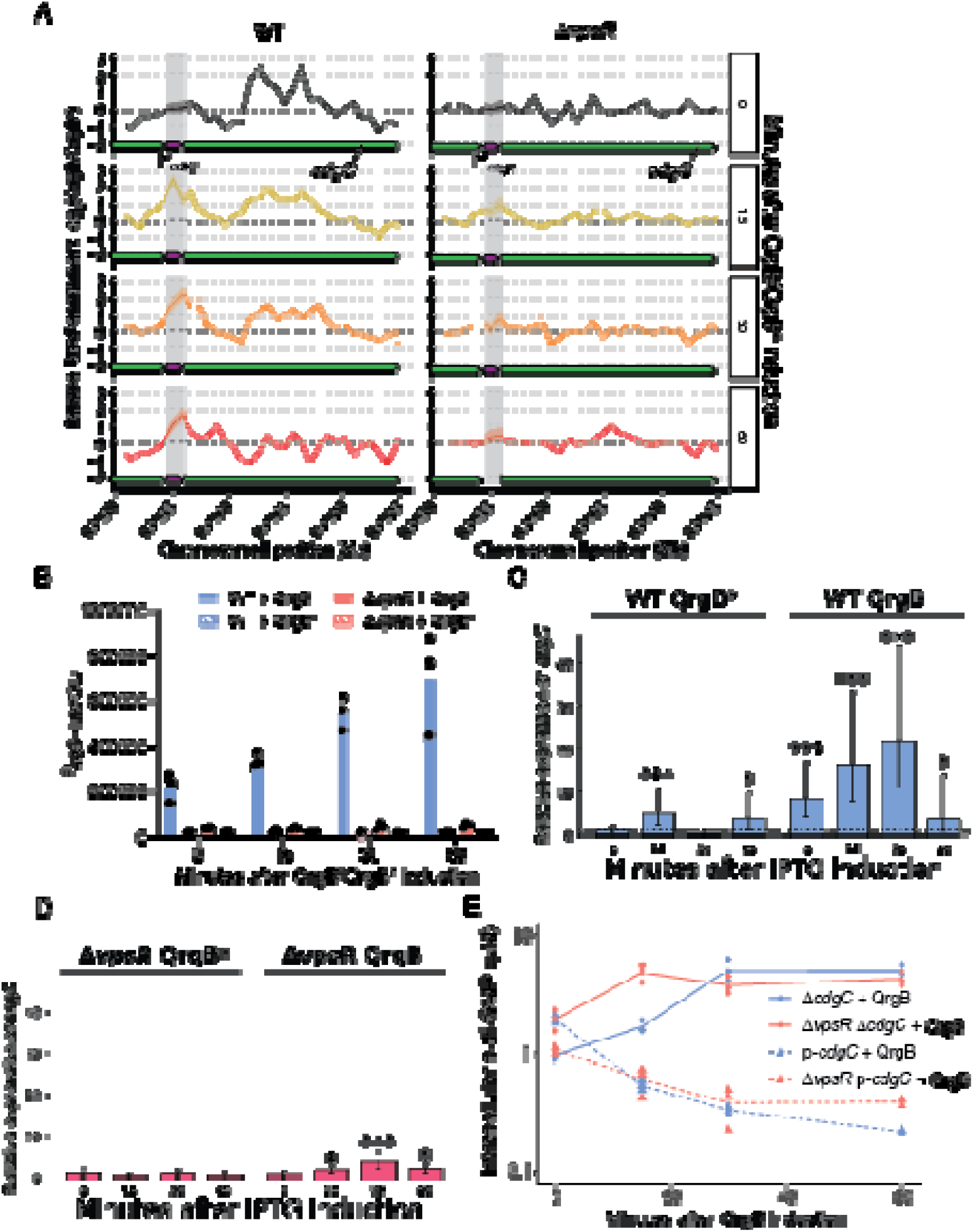
VpsR controls c-di-GMP levels through CdgC. **A)** Effect of c-di-GMP induction on RNAP occupancy at the *cdgC* locus is plotted as the mean (solid line) and 90% credible interval (shaded region). The vertical gray rectangle indicates the approximate location of the *cdgC* promoter. **B)** Expression of P*_cdgC_-lux* promoter fusion in WT and Δ*vpsR* with QrgB or QrgB* overexpression at 0, 15, 30 and 60 minutes post IPTG induction after normalization by OD_600_. Bar height represents the mean of three biological replicates; overlaid points represent the value for each of three biological replicates. **C-D)** Relative expression of *cdgC* in **C)** WT and **D)** Δ*vpsR* with QrgB or QrgB* overexpression at 0, 15, 30 and 60 minutes determined by RT-qPCR, comparing *cdgC* transcript levels (relative to a *gyrA* control) at each indicated timepoint relative to the QrgB* 0-minute timepoint in the same genetic background. Bar height and error bars indicate the mean and 95% credible intervals of a Bayesian analysis based on three biological replicates (see Methods for details). Stars indicate the posterior probability of an increase relative to baseline: *, P>0.95; **, P>0.99; ***, P>0.999. **E)** Intracellular concentrations of cdG measured by UPLC-MS/MS in Δ*cdgC*, Δ*vpsR* Δ*cdgC* mutants and *cdgC* overexpression (p-*cdgC*) in WT and Δ*vpsR* at different time points after QrgB induction. Lines are the mean value and overlaid points are the values from each of three biological replicates.

To elucidate the dynamics of c-di-GMP induction of *cdgC*, we constructed a *cdgC*-*lux* transcriptional reporter to quantify *cdgC* transcription. Consistent with our RNA-seq data, *cdgC-lux* expression increased from 0 min to 60 min after the induction of QrgB in the WT strain whereas expression in the Δ*vpsR* mutant was minimal. QrgB* expression did not induce *cdgC* transcription in either strain (**Fig 3B**). To confirm this result, we also performed RT-qPCR analysis of *cdgC* expression levels in the same samples and normalized the expression levels to QrgB* samples at 0 min. The relative expression of *cdgC* increased over time from 0 min (8-fold), 15 min (16-fold) and 30 min (22-fold) but decreased to 4-fold induction at 60 min. **(Fig. 3B**, **Fig. 3C),** likely reflecting the decrease in c-di-GMP we observed (**Fig. 1A,C**). No substantial time dependent increase in the expression of *cdgC* was seen in the Δ*vpsR* mutant (**Fig. 3D**). QrgB* expression also did not substantially induce *cdgC* transcription in either WT or Δ*vpsR* cells (with no more than 5-fold increase relative to basal expression observed even at 15-minutes), indicating this induction is a c-di-GMP specific response (**Fig. 3C and 3D**).

To test whether the increase in expression of *cdgC* was specific to QrgB or a general response to increased c-di-GMP levels, we measured *cdgC* induction upon overexpression of the known active DGC VC2224 from *V. cholerae*^26^. *cdgC* was induced 4-fold in the WT strain compared to the empty vector at 30 minutes, but no significant induction was observed in the Δ*vpsR* strain (**Fig. S3**). *cdgC* was also previously observed to be upregulated when global c-di-GMP levels are increased by several environmental or genetic conditions^11,27,28^. This indicates that *cdgC* responds to general increase in the c-di-GMP levels irrespective of the specific DGC or DGCs that are responsible for it.

### VpsR controls c-di-GMP levels through CdgC

Though CdgC contains both GGDEF and EAL domains, it lacks the catalytic residues required to produce c-di-GMP, and upon overexpression it acts as a PDE through its EAL domain to decrease c-di-GMP, thus increasing motility^29^. In addition, deletion of *cdgC* results in higher biofilm formation potentially due to loss of this PDE activity^22,30^. We therefore hypothesized that induction of *cdgC* by c-di-GMP could promote degradation of c-di-GMP in wild type and Δ*vpsT* cells, accounting for the observed negative feedback loop. Consistent with this hypothesis, expression of QrgB in a Δc*dgC* mutant resulted in high c-di-GMP levels that were not decreased at 60 minutes (**Fig. 3E**), analogous to what we observed in the Δ*vpsR* mutant (**Fig. 1A**). Conversely, overexpression of CdgC decreased the levels of c-di-GMP in Δ*vpsR* cells to levels similar to the WT (**Fig. 3E**). These results indicate that CdgC is responsible for maintaining low levels of c-di-GMP in wild type cells during the c-di-GMP response; that VpsR, independent of VpsT, induces CdgC expression to reduce c-di-GMP levels; and that CdgC expression is necessary and sufficient for VpsR-dependent feedback moderation of c-di-GMP levels.

### Identification of a counter-transcribed sRNA in *cdgC*

Visual inspection of RNA-seq fragment pileups at the *cdgC* locus qualitatively revealed the presence of an antisense RNA transcribed from the *cdgC* locus. Expression of the antisense RNA, hereafter termed <u>s</u>RNA <u>n</u>egative regulator of <u>C</u>dgC (SnrC), increased in the Δ*vpsR* mutant upon c-di-GMP induction (**Fig. S4A and S4B**). To test for the presence of SnrC and its regulation by c-di-GMP, we used RT-qPCR to quantify relative expression of *snrC* and *cdgC* in the WT and Δ*vpsR* mutant upon overexpression of QrgB, QrgB* or the empty vector after 30 min of IPTG induction. Confirming our previous observation, expression of *cdgC* was more than 10-fold higher than baseline with QrgB induction less than three-fold higher with QrgB*, and induction in the absence of *vpsR* was essentially non-existent (**Fig. S4C**). At the same time, and consistent with the RNA-seq data, *snrC* transcript levels were increased roughly 5-fold in Δ*vpsR* mutants relative to an empty vector WT control, but here there was not a substantial difference between the active and inactive QrgB **(Fig. S4D)**. Thus, the absence of VpsR appears to be the main factor permitting induction of SnrC as the density of the culture increases. We note that both the SnrC and CdgC transcripts were undetectable (Cq > 35) in our assay in a Δ*cdgC* strain (vs. Cq<31 for the corresponding *cdgC+* background in all cases), confirming the specificity of our readout.

SnrC is a counter-transcribed sRNA encoded in *cdgC*, and we thus hypothesized that SnrC could negatively regulate *cdgC* expression. To test this hypothesis, we overexpressed SnrC from a plasmid along with QrgB in wild type cells. Overexpression of SnrC increased the levels of c-di-GMP, potentially by counteracting *cdgC* expression (**Fig. S4E**). This increase was specific to SnrC as expression of RpoA RNA and the complement of SnrC (sense-*snrC*) did not increase c-di-GMP (**Fig. S4E**). c-di-GMP levels were high in a Δ*cdgC* mutant and unaffected by SnrC, sense-*snrC*, or RpoA RNA, indicating that the overexpression of SnrC increases c-di-GMP levels specifically through CdgC (**Fig. S4F**). These results indicate the presence of a counter-transcribed RNA in *cdgC* that reduces *cdgC* expression post-transcriptionally. However, the precise mechanism of regulation of *snrC* by VpsR, the role (if any) of c-di-GMP in that regulation, and the physiological role of *SnrC* in modulating c-di-GMP concentrations requires further study. A concomitant increase in RNA polymerase occupancy near the likely *snrC* transcription start site can be seen at the 60-minute timepoint in the Δ*vpsR* strain (**Fig. 3A**; position 0.7965 Mbp), although the occupancy is relatively weak likely due to the fairly low transcript levels seen for *SnrC* under this condition.

### Negative feedback by CdgC promotes growth during QrgB induction

Elevated intracellular c-di-GMP levels exert a detrimental impact on the growth of *V. cholerae*^26^. We hypothesized that the sustained high levels of c-di-GMP caused by the absence of CdgC or VpsR would be especially disadvantageous for *V. cholerae* growth. We thus assessed the growth of wild type, Δ*vpsR*, Δ*cdgC*, and Δ*cdgC/*Δ*vpsR* mutants while inducing QrgB with increasing IPTG concentrations. Notably, at the highest IPTG concentrations tested (0.625 and 1.25 mM) the Δ*cdgC*, Δ*vpsR*, and Δ*vpsR* Δ*cdgC* mutants exhibited reduced growth compared to wild type. In contrast, at the lower IPTG concentrations (0.0 and 0.15 mM), no discernible difference in growth emerged between wild type and mutants, implying that constitutively elevated c-di-GMP levels curtail *V. cholerae* growth and such levels of c-di-GMP are controlled by VpsR and CdgC (**Fig. 4A**). Our growth curves also further affirmed the epistatic nature of VpsR and CdgC, as the Δ*vpsR* Δ*cdgC* double mutant displayed practically identical growth to the Δ*vpsR* mutant even at the highest tested concentration of IPTG (**Fig. 4A**, right panel). As noted above, each of these strains are in a Δ*vpsL* mutant background, ruling out the possibility that the observed growth defects are due to biofilm formation itself or an artifact of cellular aggregation.

**Figure 4.**
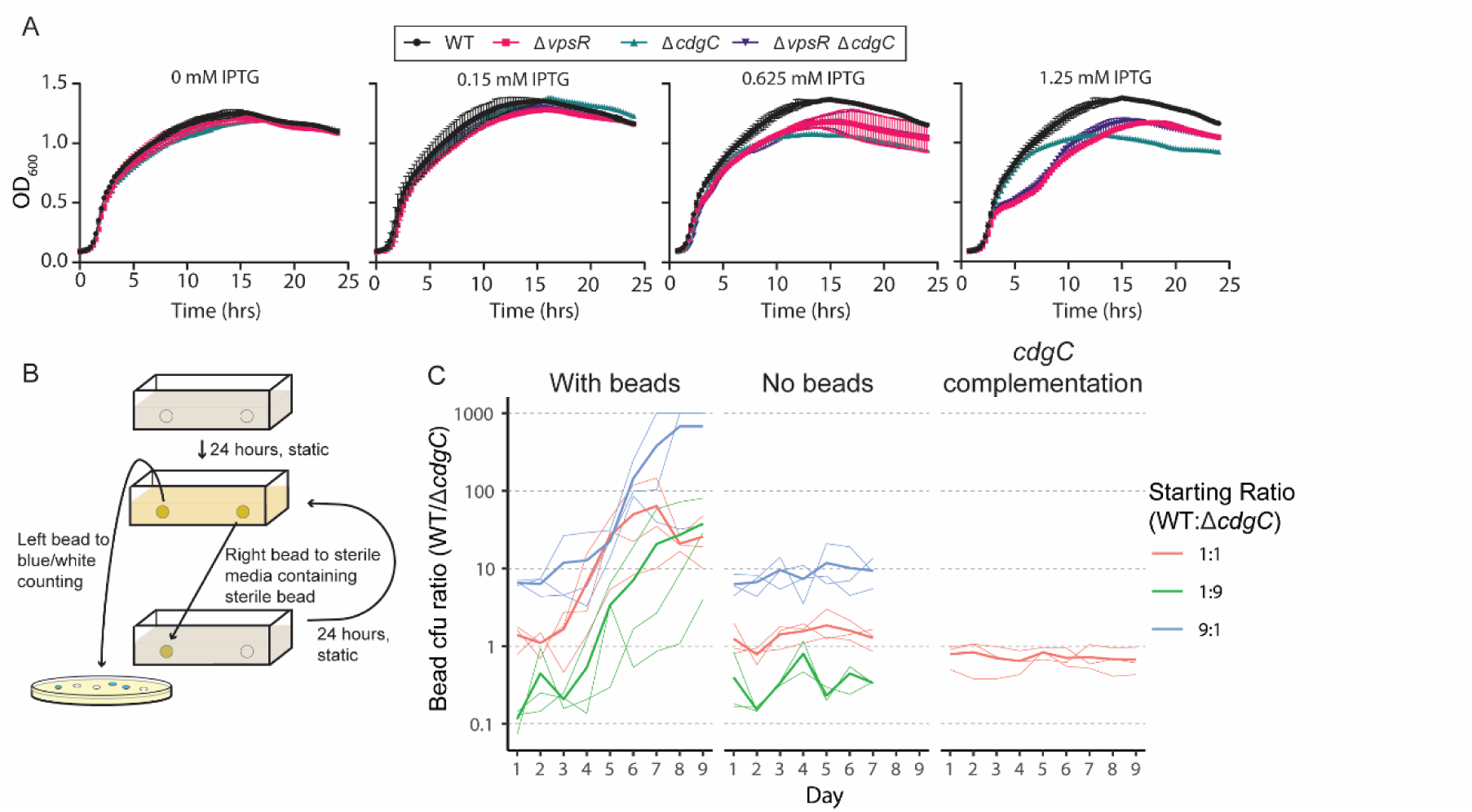
CdgC promotes balanced biofilm formation and dispersal. **A)** Growth curves of WT, Δ*vpsR*, Δ*cdgC*, and Δ*vpsR* Δ*cdgC* strains containing QrgB induced with 0, 0.15, 0.625 and 1.25 mM IPTG. Data represent the mean ± SD of 3 biological replicates. **B)** Schematic demonstrating competition assay. Biofilm-coated beads are either used to count blue/white colonies or to inoculate sterile medium containing a fresh sterile bead every 24 hours. Static growth requires cells in the biofilm to disperse from the left bead and form a new biofilm on the right bead to be competitive in the assay. **C)** Data showing ratio of colony forming units (cfu) between WT (*lac-*) and Δ*cdgC* (*lac+*) in the presence (left plot) and absence (middle plot) of beads and WT (*lac+*); or Δ*cdgC lacZ::*P*_cdgC_*:*cdgC (lac-)* in the presence of beads (right plot). Thin lines represent independent biological replicates, and thick lines represent the mean ratio of three biological replicates for each starting ratio. The ratios were clipped to a maximum value of 1,000.

### CdgC allows balanced biofilm formation and dispersal behaviors to enhance fitness in fluctuating environments

High and low c-di-GMP levels drive attachment and dispersal, respectively. Following dispersal, bacteria may colonize new surfaces. Both biofilm formation and dispersal are crucial for survival, and for successful dissemination of *V. cholerae* during human infection and colonization in aquatic reservoirs^4,31^. Prior work demonstrated that biofilm formation is increased in a *cdgC* mutant^29^. Although loss of CdgC has been associated with increased biofilm formation, we hypothesized that Δ*cdgC* cells could have a fitness cost in physiologically relevant environments due to decreased dispersal from biofilms owing to dysregulated c-di-GMP levels. That is to say, we predict that cells lacking CdgC will over-commit to entering, and subsequently remain in, a biofilm when c-di-GMP levels increase. We therefore predicted that loss of the c-di-GMP negative feedback loop in Δ*cdgC* cells should lead to decreased fitness in a fluctuating environment requiring well-balanced biofilm formation/dispersal dynamics.

To test whether CdgC promoted fitness in fluctuating environments through balancing biofilm formation and dispersal, we performed a competition assay modeled after biofilm evolution experiments created by the Cooper laboratory^32,33^. We competed wild type *lac*^−^ strains against Δ*cdgC lac*^+^ strains based on their ability to disperse from biofilms coating a polystyrene bead to form a new biofilm on fresh beads. The competition assay was performed by serially passaging biofilm-coated beads into sterile media containing a fresh sterile bead every 24 hours (**Fig. 4B**). These experiments were performed in a *vpsL^+^* background to allow normal biofilm formation, and the QrgB expression plasmids were not used, allowing us to investigate the behavior of the cells in the context of their native c-di-GMP regulatory network. The competition/transfer process was continued for 9 days, and each day the bead that was not passaged to fresh media was vortexed in fresh solution and plated, and the ratio of wild type colonies (white on X-gal-containing plates) to Δ*cdgC* colonies (blue on X-gal-containing plates) was calculated by comparing the colony forming units (CFUs) of each color. As shown in **Fig. 4C**, regardless of the starting ratio of wild type vs. Δ*cdgC* cells, wild type was able to outcompete Δ*cdgC*, potentially due to poor dispersal of cells lacking CdgC.

As a control we competed the wild type *lac*^−^ strains and Δ*cdgC lac*^+^ strains in equivalent media without beads, which showed no change in the ratio of wild type to Δ*cdgC* cells over time, indicating there is no difference in the growth rate of wild type and *cdgC* mutant cells when grown planktonically in minimal media (**Fig. 4C**). We also complemented the *cdgC* gene along with its native promoter region by placing them in the *lacZ* locus in the Δ*cdgC* mutant. After complementing *cdgC* under control of its native promoter, there was no difference in fitness between wild type *lac*^+^ cells and Δ*cdgC lacZ*::P*_cdgC_*:*cgdC* cells (**Fig. 4C**). This indicates that the *cdgC* gene *per se* is responsible for loss of fitness in our competition assay, effectively ruling out any possibility that deletion of *cdgC* caused other effects. Taking together, these data suggest that the PDE activity of CdgC is responsible for limiting the levels of c-di-GMP priming the population of cells to efficiently switch between biofilm formation and dispersal, which increases the fitness of *V. cholerae* in dynamic environments.

## Discussion

Both biofilm formation and dispersal are important for the successful colonization and dispersal of *V. cholerae* in the human gut and in other environments^4,34^. Biofilms themselves have heterogenous c-di-GMP levels, exhibiting higher c-di-GMP levels in the core of the biofilm compared to the edges; this heterogeneity enables dispersal and increased fitness in fluctuating environments^35^. Hyper biofilm forming mutants possessing high intracellular c-di-GMP display decreased colonization of mice when compared to wild type, potentially due to poor dispersal^36^. Moreover, high levels of c-di-GMP in *V. fischeri* impair squid colonization^37^. In this work, we identified a regulatory circuit that influences the balance of *V. cholerae* biofilm formation and dispersal in response to c-di-GMP induction.

Here, we show that through VpsR, which is the master regulator of the biofilm response to c-di-GMP, the PDE CdgC enforces negative feedback on c-di-GMP levels. Previous findings have demonstrated CdgC to be differentially regulated in a *vpsR* mutant, the *cdgC* promoter to possess a predicted VpsR binding site, and *cdgC* expression was upregulated during increased global c-di-GMP levels in several conditions^11,14,27,28^. We demonstrated that the c-di-GMP → VpsR → CdgC ⊣ c-di-GMP regulatory path (**Fig. 5**) serves as a pulse generator, bringing about a short-duration rise in c-di-GMP levels after c-di-GMP induction by an external cue. This property of the c-di-GMP → VpsR → CdgC negative feedback loop is apparent both from our LC-MS analysis of c-di-GMP levels in wild type vs. Δ*vpsR* cells, and by our RNA-seq analysis of flagellar genes exhibiting a pulse of repression in wild type and Δ*vpsT* cells, but not in Δ*vpsR* cells (**Fig. 1**). We additionally showed that induction of *cdgC* by VpsR provides increased fitness to *V. cholerae* when biofilm formation and dispersal must exist in balance with each other (**Fig. 4**).

**Figure 5.**
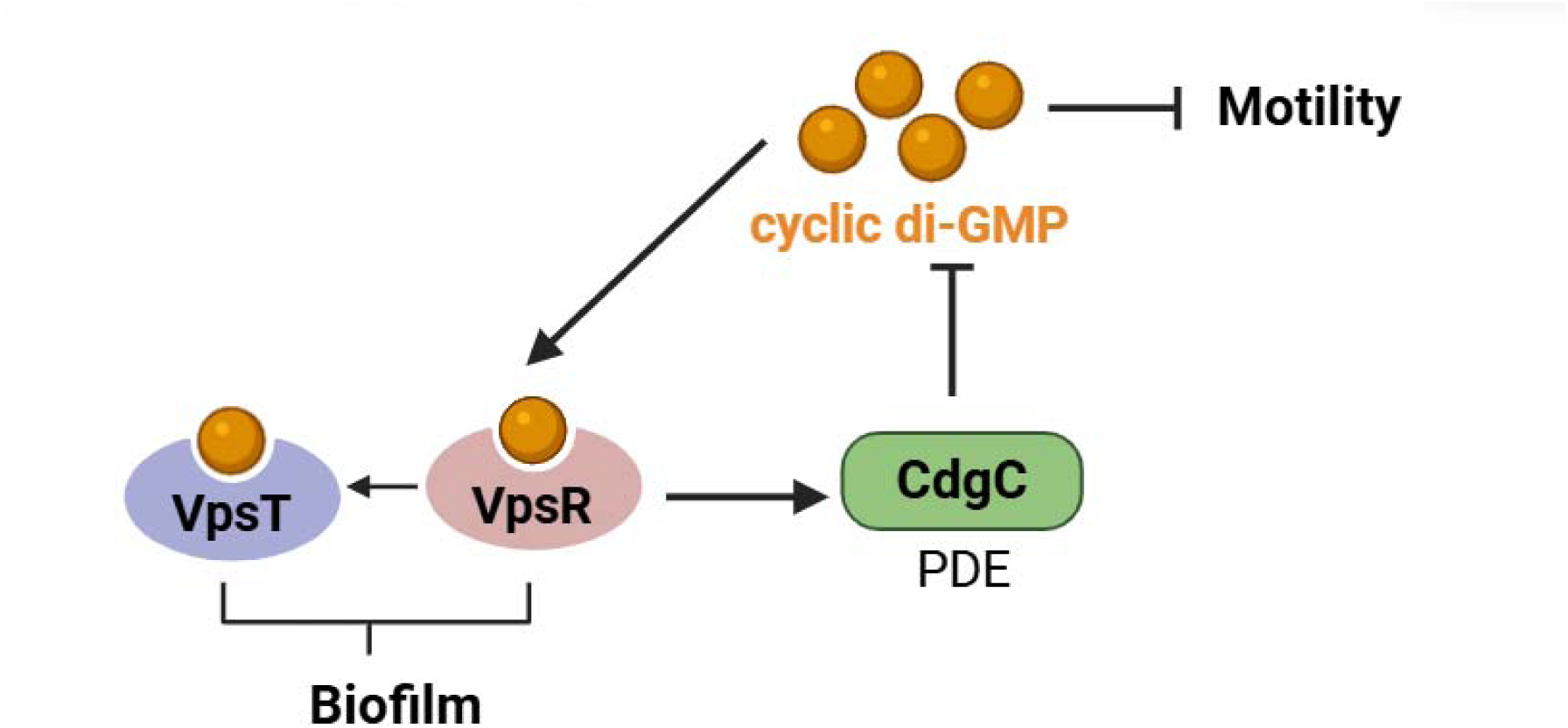
Model for feedback regulation of c-di-GMP levels by CdgC via VpsR. When intracellular levels of c-di-GMP increase, VpsR in the presence of c-di-GMP activates expression of CdgC which decreases c-di-GMP levels via its PDE activity. This feedback enables *V. cholerae* to quickly switch between motile and sessile lifestyles without becoming over-committed to a biofilm lifestyle.

Positive feedback on c-di-GMP levels was recently proposed to occur through VpsT-dependent activation of the DGC VpvC^38^. However, we did not detect evidence for induction of VpvC expression under the conditions of our experiments (**Fig. 2A**, locus tag VC2454). We find it likely that differing laboratory conditions may explain our identification of a negative feedback loop whereas others have posited that positive feedback occurs; the two may both exist, but act under differing environmental conditions. Indeed, the potential existence of opposing feedback loops on c-di-GMP that can be turned on or off depending on environmental cues is intriguing, and it would underscore the benefits of balancing c-di-GMP levels until the proper signals are in place to fully commit to motility, biofilm production, or dispersal. Alternatively, our study used *V. cholerae* strain C6706 while ^38^ utilized strain E7946, which could also be responsible for the different results.

Global analysis of regulators in *Escherichia coli* revealed that several stress response regulators including the c-di-GMP phosphodiesterase PdeH and the biofilm regulator CsgD are readily degraded^39^. Here, we have identified a putative countertranscribed sRNA, SnrC, that counteracts the expression of *cdgC,* adding another layer of regulation for this key PDE. Although overexpression of *snrC* counteracts the expression of *cdgC,* thereby increasing c-di-GMP levels, the conditions at which it is expressed are still unclear. SnrC may be induced by specific environmental conditions to counteract *cdgC* expression to fine-tune the level of c-di-GMP for biofilm formation independent of VpsR regulation at the level of transcription. CdgCs EAL activity could also be modulated by its degenerate GGDEF domain and N-terminal sensory domain. Thus, our work and others suggest that CdgC is a central modulator of c-di-GMP in *V. cholerae*, while indicating that we still do not yet fully understand the range of upstream regulatory inputs acting on it.

Some clues as to the as-yet unidentified additional modulators of CdgC can be derived via structural bioinformatic approaches. For example, we note that the degenerate GGDEF domain of CdgC may bind c-di-GMP to modulate PDE activity. At the N-terminus, the first 156 residues of the structural prediction of CdgC (AFDB model AF-Q9KLF9-F1) model the CdgC sensing domain with high confidence^40,41^. Structural alignment of the sensing domain against structures in the Protein Data Bank using Foldseek^42^ (using the Foldseek webserver with default settings) reveals that the sensing domain of CdgC is similar to light-sensing GAF domains found on the DGC Dcsbis from *Pseudomonas aeruginosa*^43^ (E-value 2.37*10^−10^) and Cph2 from *Synechocystis* sp. PCC 6803 (E-value 5.37*10^−7^), the DGC and EAL activity of which is likely to be modulated by light sensed by the GAF domains in Cph2^44^. CdgC PDE activity may be modulated by light, providing additional control of its output (and perhaps altering the activity of CdgC in a mammalian host vs. a planktonic environment or on the surface of an arthropod).

Understanding how the genetic circuits in the c-di-GMP metabolic network identified here impact various aspects of *V. cholerae* pathogenesis will require further investigation of both the transcriptional regulators (like VpsR) and post-transcriptional regulators (like SnrC) that may modulate CdgC activity. In addition, while we demonstrated here that a proper balance of biofilm formation and dispersal through VpsR and CdgC enhances fitness, it will be useful to understand precisely when CdgC and other regulators of *cdgC* play their roles in c-di-GMP metabolism.

## Supporting information

Response to Reviewers

## Supplemental Figures

**Figure S1.**
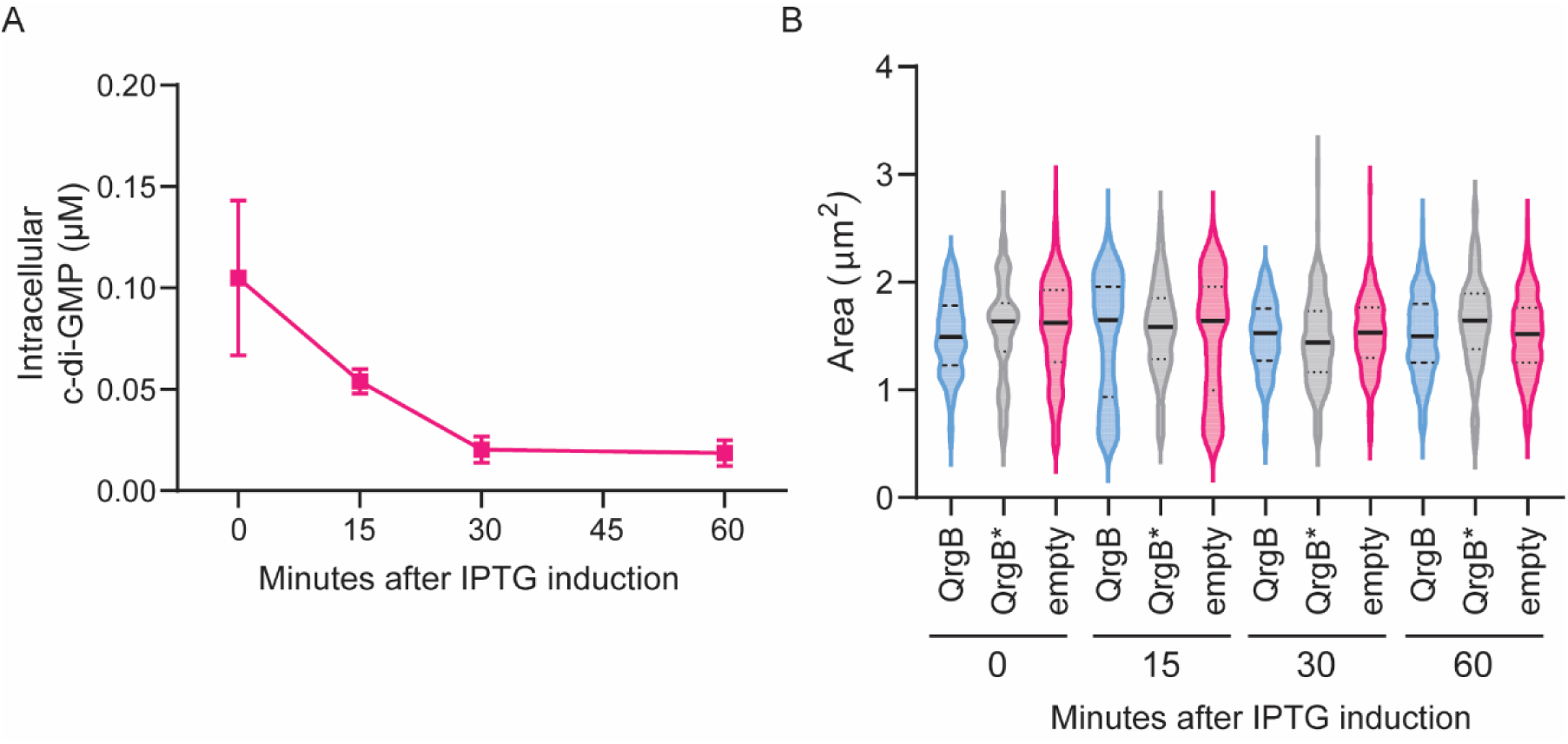
**A)** Intracellular concentrations of c-di-GMP measured by UPLC-MS/MS in WT with an empty vector induced with IPTG at an OD of 1 at 0-minute timepoint. Data represents the mean ± SEM. from 3 biological replicates. **B)** Violin plot representing area of the WT cells expressing QrgB, QrgB* or an empty vector measured from single cell phase contrast microscopy images. Randomly selected 200 representative cells were analyzed from 3 biological replicates. The solid line represents median and the dotted lines represent the upper and lower quartiles.

**Figure S2.**
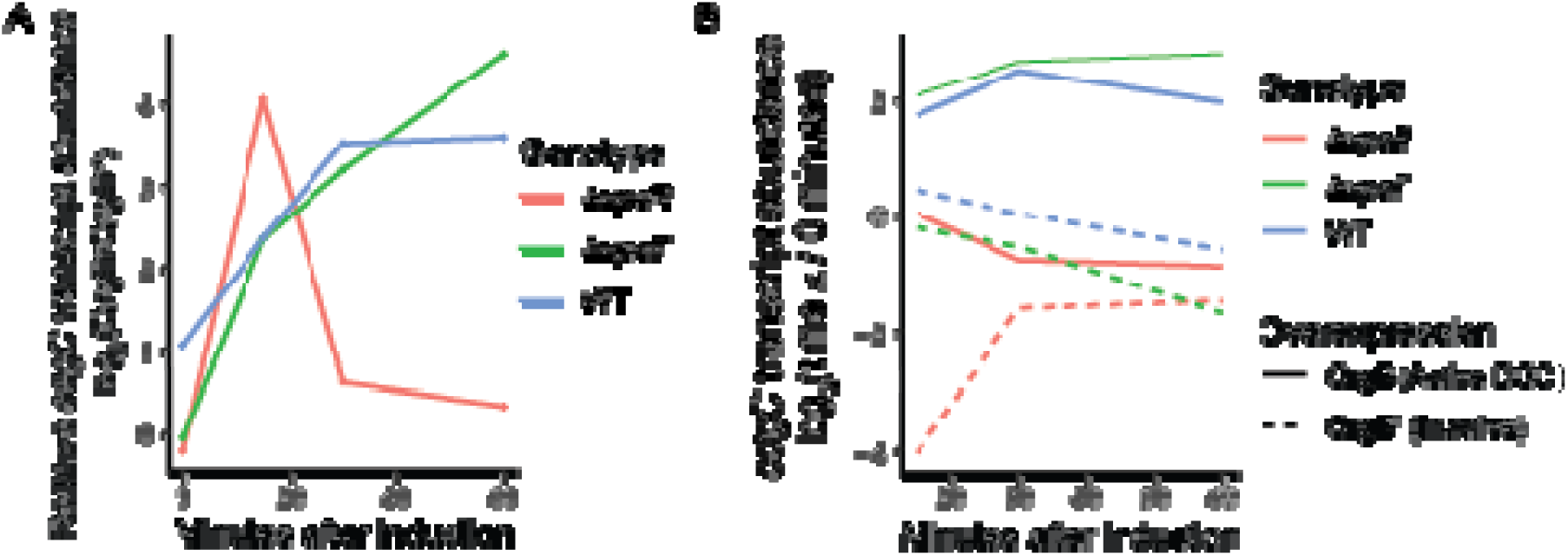
**A)** Analysis of RNA-seq data expressed as relative *cdgC* transcript abundance with QrgB* at each timepoint and genotype as the baseline for comparison. Increased *cdgC* expression over time is evident in WT and Δ*vpsT*. **B)** Analysis of RNA-seq data expressed as relative *cdgC* transcript abundance with 0 minutes after QrgB or QrgB* induction as the baseline for comparison. Induction of *cdgC* expression in wild type and Δ*vpsT* cells overexpressing QrgB is evident.

**Figure S3:**
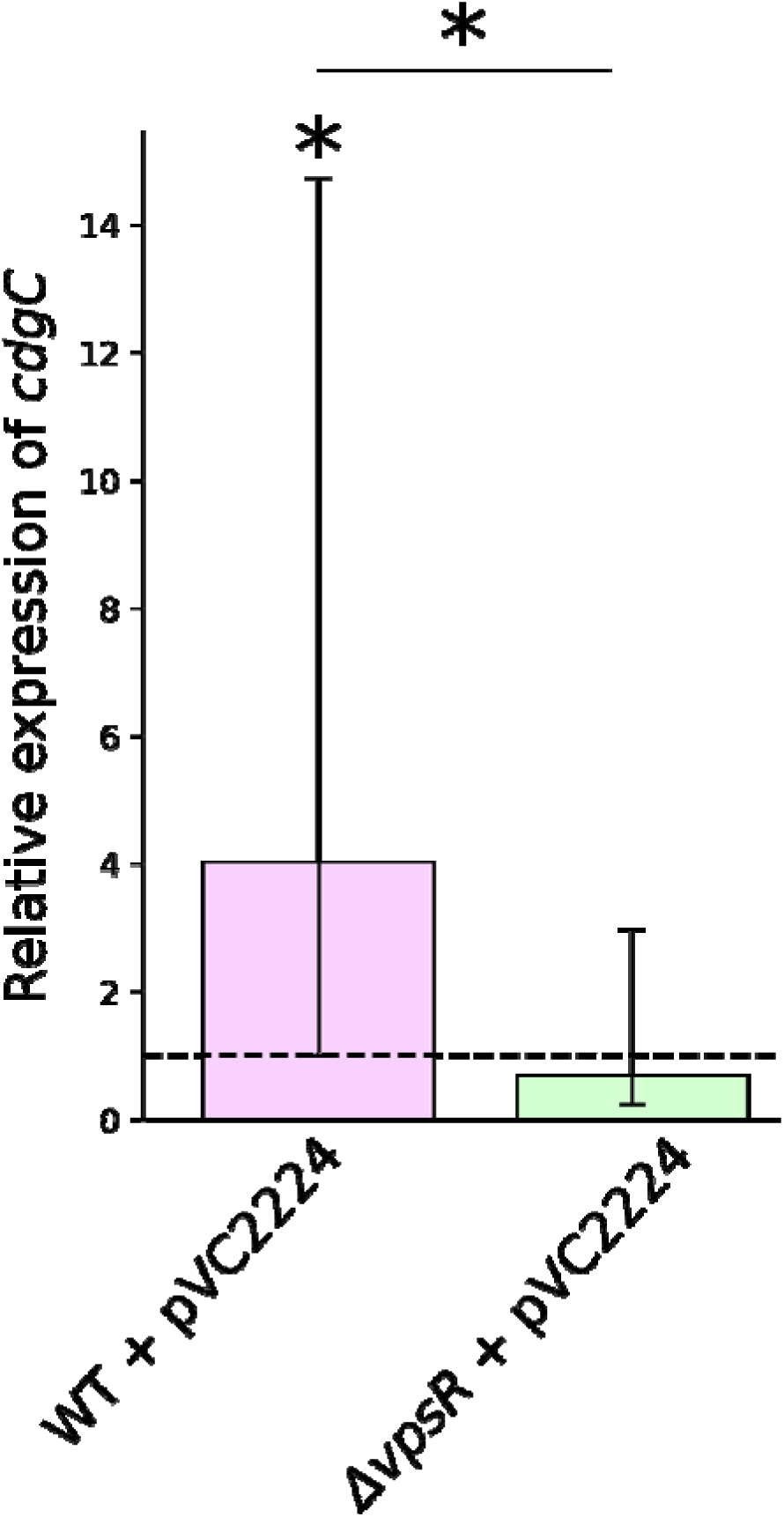
Relative expression of *cdgC* (compared with a *rpoC* reference) in WT with VC2224 or Δ*vpsR* with VC2224, relative to WT with empty vector, assessed via RT-qPCR. Bar height and error bars indicate the mean and 95% credible intervals of a Bayesian analysis based on two (WT, pVC2224) or three biological replicates (see Methods for details). Stars indicate the posterior probability of an increase relative to baseline (or for the indicated comparison): *, P>0.95; **, P>0.99; ***, P>0.999.

**Figure S4:**
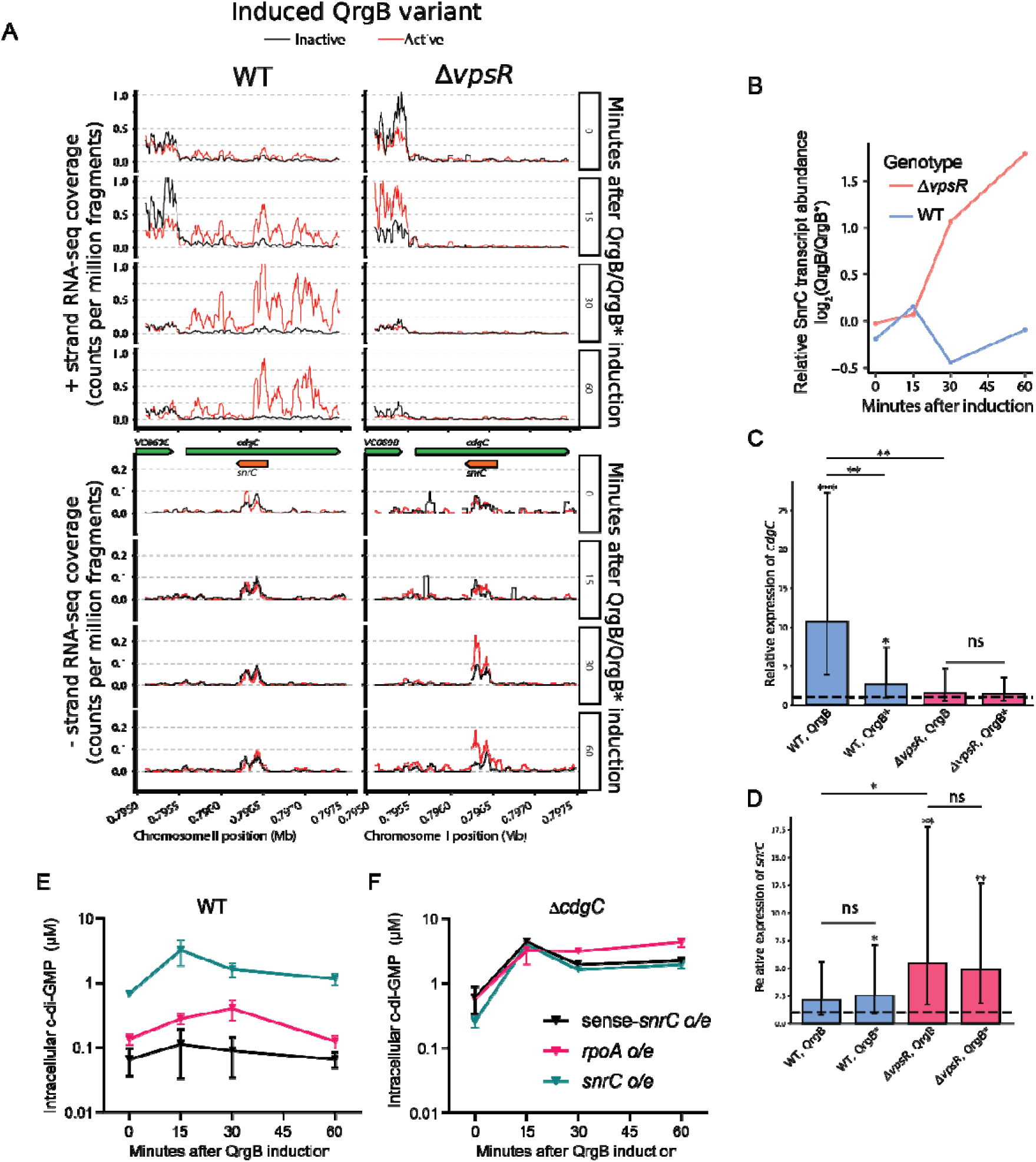
**A)** RNA-seq fragment abundance for the plus (top) and minus (bottom) strand are plotted in the vicinity of *cdgC/snrC* for WT (left) and Δ*vpsR* (right), for each of QrgB (Active) and QrgB* (Inactive) induction at each timepoint. Note y-axis limits differ for the plus vs. minus strand due to stronger sense transcription in this region. **B)** Effect of c-di-GMP induction on expression of *snrC* (measure via RNA-seq) is plotted at each timepoint. **C)** Expression of *cdgC* in WT and Δ*vpsR* (via RT-qPCR, vs. a *rpoC* reference) with QrgB or QrgB* induction (as indicated) relative to empty vector, after 30 minutes of induction. Bar height and error bars indicate the posterior mean and 95% credible intervals of a Bayesian analysis based on three biological replicates (see Methods for details). Stars indicate the posterior probability of an increase relative to baseline (or for the indicated comparison): *, P>0.95; **, P>0.99; ***, P>0.999. **D)** As in panel C, for *snrC* expression relative to *rpoC.* **E-F)** Intracellular concentrations of c-di-GMP measured by UPLC-MS/MS in **E)** WT and **F)** Δ*cdgC* mutant with ectopic overexpression of *snrC*, and control transcripts RpoA and the reverse complement of *snrC* (sense-*snrC*). Data in E and F represent the mean ± SEM. from 3 biological replicates.

## Methods

### Growth conditions and gene manipulations

All the strains are derivatives of *V. cholerae* O1 El Tor biotype strain C6706str2. The list of strains, plasmids and oligonucleotides used in this study are listed in supplementary table S1-S3. CW2034 (Δ*vpsL*) is designated as “wild type” in all the experiments and all other strains were derived from this background except for those used for the biofilm assays in Fig. 4C, for which wild-type C6706str2 (*vpsL+*) was used. The strains were grown in LB broth at 37°C with shaking. Antibiotics were used at the following concentrations; kanamycin (50 μg/ml) and ampicillin (50 μg/ml). 1 mM IPTG was used for induction of QrgB of QrgB*. Gene deletions and insertions were performed using homologous recombination and the pKAS32 suicide plasmid as described previously^46^. Overexpression plasmids were constructed in the P*_tac_* inducible vector pEVS143^47,48^ or pMMB67EH^46^ using PCR and Gibson assembly and verified by sequencing.

### Quantification of intracellular c-di-GMP levels

C-di-GMP quantification was performed using UPLC-MS/MS as described previously^26^. The cells were grown to OD_600_ of 1, after which 1 mM IPTG was added and samples were taken at 0, 15, 30 and 60 minutes after the addition of IPTG. Briefly 1 ml of cultures were spun down at 20,000 r.c.f for 2 min, resuspended with 100 µl nucleotide extraction solution (40% acetonitrile, 40% methanol, 0.1% formic acid, and 19.9% water) and incubated at -20°C for 20 minutes. Following this, the samples were centrifuged at 20,000 r.c.f for 15 minutes and the supernatant was transferred to a fresh tube and dried under vacuum. The samples are then resuspended in 100 µl HPLC grade water, and 10 µl of this solution was injected into the UPLC-MS/MS system (Waters, Xevo).

Electrospray ionization using multiple reaction monitoring in negative-ion mode at m/z 689.16→344.31 was used to detect cdG. The MS parameters were as follows: capillary voltage, 3.5 kV; cone voltage, 50 V; collision energy, 34 V; source temperature, 110 °C; desolvation temperature, 350 °C; cone gas flow (nitrogen), 50 L/h; desolvation gas flow (nitrogen), 800 L/h; collision gas flow (nitrogen), 0.15 mL/min; and multiplier voltage, 650 V. A Waters BEH C18 2.1 × 50 mm column with a flow rate of 0.3 mL/min with the following gradient was used: solvent A (10 mM tributylamine plus 15 mM acetic acid in 97:3 water:methanol) to solvent B (methanol t = 0 min; A-90%:B-10%, t = 2.5 min; A-80%:B-20%, t = 3 min; A-80%:B-20%, t=6.0 min; A-35%:B-65%, t = 6.5 min; A-5%:B-95%, t = 8.0 min; A-5%:B-95%, t = 8.01 min; A-90%:B-10%, t = 10 min; A-90%:B-10% (end of gradient). 125, 62.5, 31.25, 15.62, 7.81, 3.625, 1.9 nM of c-di-GMP (Biolog) was used to generate a standard curve for calculating the c-di-GMP concentrations in the sample. The intracellular concentration was determined by dividing the total moles of c-di-GMP by the intracellular volume of bacteria. The intracellular volume was determined by multiplying the number of CFU in each sample by the volume of one bacterium which was previously determined to be 6.46 × 10^−16^ L, and which did not show meaningful variations over our induction time courses (**Fig. S1B**).

### CdgC Promoter-*lux* analysis

The promoter of *cdgC* was cloned in pBBR-lux reporter^49^ and conjugated into wild-type and Δ*vpsR* cells with QrgB and QrgB*. Overnight cultures were diluted 1:1000 in flasks and grown in triplicates with Ampicillin (50 µg/ml) and Chloramphenicol (10 µg/ml) to OD_600_ of 1 at which IPTG was added to the final concentration of 1 mM. 100 μl samples were taken for measurements in Costar 96-well black bottom plates at 0, 15, 30 and 60 minutes after IPTG induction and the respective OD_600_ of the samples were measured. Luminescence was measured using Envision Multi-label plate reader (Perkin-Elmer) and the luminescence was normalized to OD_600_.

### RT-qPCR analysis

For RT-qPCR analysis of figure 3C and 3D, 1 mL of cultures from *cdgC*-lux at 0 min, 15 min, 30 min and 60 min was pelleted. RNA was isolated using TRIzol reagent (Thermo Fisher). 5 µg of RNA was treated with Turbo DNase (Thermo Fisher) and cDNA synthesis was performed using Superscript III reverse transcriptase (Thermo Fisher). 0.625 µM of primers AAR195 and AAR196 with 0.78 ng of cDNA and 2x SYBR Green Master Mix (Applied Biosystem) in a 25 µL reaction. The reactions were performed in technical triplicates from each of three biological replicates. RT–qPCR reaction conditions are 95 °C for 20 s, then 40 cycles of 95 °C for 2 s and 60 °C for 30 s in the QuantStudio 3 Real-Time PCR system (Applied Biosystems). *gyrA* was used as an endogenous reference gene for relative quantification (Δ*C*_t_). For the RT-qPCR analysis in Figure S4C and S4D, purified nucleic acid was treated twice with TURBO DNAse in the following manner. To 45 uL of purified nucleic acid, 5 µL of 10X TURBO DNAse buffer was added. After mixing thoroughly, 1 µL of TURBO DNAse was added, mixed, and incubated at 37°C for 1 hour. An additional 1 µL of TURBO DNAse was added, mixed, and incubated at 37°C for 1 hour. RNA was purified using a Zymo RNA Clean and Concentrate kit. RNA was eluted from spin columns using 20 µL of nuclease-free water. RNA concentration was measured via nanodrop. For first strand cDNA synthesis we used ProtoScript II First Strand cDNA Synthesis Kit (NEB, #E6560). 20 µL reverse transcription reactions were prepared using 100 ng RNA. Biotinylated reverse transcriptase primer oJS0093, oJS0094, or oJS0095 was used at a final concentration of µM. A separate reaction with no primer was included for every RNA template as a control. 10 µL of 2X Protoscript II reaction mix was added and the solution was mixed. 2 µL of Protoscript II enzyme mix and sufficient water to bring the final volume to 20 µL, and the solution was mixed. Reactions were performed at 48° C for one hour, followed by heat inactivation at 80°C.

Biotin-linked cDNA was pulled down using the following protocol. 2 mL of Dynabeads MyOne Straptavidin C1 magnetic bead slurry was placed on a magnetic rack and the supernatent was discarded. Beads were resuspended in 1 mL of binding buffer (5 mM Tris-HCl, pH 7.5, 0.5 mM EDTA, 1 M NaCl, 0.05% w/v Tween-20) and placed on a magnetic rack. Supernatant was discarded. For a total of three rounds of washes with mL of binding buffer, beads were resuspended in 0.2 mL of binding buffer, placed on a magnetic rack, and supernatant was discarded. Beads were resuspended in 1 mL of blocking buffer (binding buffer supplemented with 1 mg/mL ssDNA from salmon testes, catalog number D7656, Sigma Aldrich). Beads were pre-blocked in a blocking buffer for 15 minutes at room temperature with constant rotation. Beads were placed on a magnetic rack and supernatant was discarded. Pre-blocked beads were resuspended in 2 mL of 2x binding buffer.

20 µL of beads were distributed to each tube and an entire 20 µL, heat-inactivated reverse transcription reaction was added to bead slurry and rotated continuously at room temperature for 15 minutes. Tubes were placed on a magnetic rack and the supernatant was discarded. Beads were washed twice with 100 µL of wash buffer (5 mM Tris-HCl, pH 7.5, 0.5 mM EDTA, 1 M NaCl, 2% w/v Tween-20). Beads were resuspended in 100 µL of wash buffer and the slurry was transferred to a fresh PCR tube. The tube was placed on a magnetic rack, and the supernatant was discarded. Beads were washed once more with 100 µL wash buffer, placed on a magnet, and supernatant discarded. Beads were resuspended in 100 µL Tris-buffered saline (10 mM Tris-HCl, pH 7.5, 150 mM NaCl), placed on a magnet, and the supernatant was discarded. To elute cDNA from beads, 20 µL pure water was added and slurry was heated to 95C in a thermocycler. Tubes were immediately transferred to a magnetic rack, and 2 µL of supernatant was added to qPCR reactions.

qPCR reactions were performed at 20 µL final volumes using 10 µL of SuperUp 2x master mix, 2 µL of cDNA eluate, 2 µL of each pre-mixed primer pair (oJS0035 and oJS0036, oJS0025 and oJS0026, or oJS0029 and oJS0030), and 6 µL of water. For additional controls for primer (cDNA primer and qPCR primer) specificities, we set up qPCR reactions for every qPCR primer pair with every cDNA. No negative controls produced detectable PCR products within 45 cycles. *rpoC* was used as an endogenous reference gene for relative quantification (Δ*C*_t_). Relative fold expression was calculated using 2^−ΔΔCt^method^50^ with a Bayesian mixed-effects model applied to calculate posterior inferences regarding the expression levels. We fitted the models using the brms package in R, with fixed effects for primer, biological condition (i.e., timepoint and/or genotype), and primer:condition interactions, and random effects for each separate biological sample (and, in the case of Fig. 3, for each technical replicate:primer combination). Fits were performed in the Ct space, using a uniform(0,40) prior on the primer effects, normal(0,4) primer on all other fixed effects, and a Student t-distribution prior with 3 degrees of freedom and a scale parameter of 1 for the random effects.

Residuals were modeled using a Student t-distribution with default priors. Convergence was assessed via Rhat and posterior predictive plotting.

### Microscopy

To assess cell volumes across growth conditions, WT CW2034 cells were grown with expression plasmids for QrgB, QrgB* or an empty vector in three biological replicates to OD_600_ of 1, 1 mM IPTG was added and samples were taken at 0, 15, 30 and 60 minutes after the addition of IPTG. 2 μl of each culture were dropped on an 1% agarose pad in a glass slide and was covered with a glass cover slip. Cells were imaged using a Leica DMI 6000 microscope using phase contrast with an HC PL APO 63x/1.4 numeric aperture oil Ph3 CS2 objective. Images were captured with an Orca-ER digital camera (Hamamatsu) controlled by Leica Application Suite X (Leica). Areas of the cells were calculated using Image J software^51^.

### Growth curve assays

Overnight cultures were diluted 1:500 into 150 μL LB supplemented with ampicillin (100 µg/ml) with 0 mM, 0.125 mM, 0.625 mM or 1.25 mM IPTG for the induction of QrgB. Growth was determined by measuring OD_600_ every 15 minutes for 24 hours at 35°C with shaking using an Infinite F500 Microplate Reader (Tecan).

### Biofilm competition assay

The biofilm dispersal assay was performed as described previously [29] with modifications as indicated below. Wild type *lacZ^−^* and Δ*cdgC lacZ*^+^ or wild type *lacZ*^+^ strain and Δ*cdgC lacZ::*P_cdgC_-*cdgC* strains were used for biofilm competition assays. The cells were inoculated in the ratio of 1:1, 9:1 or 1:9 to OD_600_ 0.05 in 10 ml of minimal media (1x M9 salts, 0.4% glucose, 2 mM MgSO_4_, 100 μM CaCl_2_) containing polymyxin B (40 μg/mL) in 4 well sterile plates (127.8 x 85.5 mm, Thermo Scientific). *V. cholerae* C6706 is naturally resistant to polymyxin B, and it was included to minimize contamination. Biofilm competition assay was performed as described in Figure 4B. Two polystyrene beads of 7 mm diameter (Costar) were placed 3 cm apart in a single well and statically incubated at 35°C for 24 hours. After 24 hours, the bead on the left side was washed in PBS by gently shaking the beads in 8 mL PBS in a petri dish (60 x 15 mm) for 1 minute, transferred to a 2 mL tube, and vortexed with 1 mL LB for 2 minutes. Dilutions of the resulting suspension were plated on LB agar containing polymyxin B with X-gal and IPTG and subsequently scored for blue and white colonies. The bead on the right was also washed with PBS similarly and placed into a new plate on the left side containing fresh minimal media with polymyxin B and a fresh bead was placed 3 cm away on the right side. After 24 hours the same process was repeated, following the same cycle for a total of 9 days. For control experiments a similar setup was used without the beads.

### Cell growth and crosslinking for RNA-seq and RNAP ChIP-seq

For each biological replicate of each genotype, cells were grown in LB supplemented with a final concentration of 100 μg/ml ampicillin. Overnight cultures were diluted to an OD_600_ = 0.05 in 500 mL of LB pre-warmed to 35°C containing 100 μg/ml ampicillin.

Cultures were incubated at 35°C with constant shaking at 200 rpm and cultured to an OD_600_ = 1.0, at which time the 0 minute timepoint aliquots were rapidly collected as follows: 200 μL of culture was mixed with 1 mL of DNA/RNA shield (Zymo Research Corporation) to be used for RNA extraction (described later) and 100 mL of culture was aliquoted to a pre-warmed 250 mL flask and treated with 75 μg/mL rifampicin.

Immediately following initial time 0 sample collection, IPTG was added to the remaining 400 mL of original culture to a final concentration of 1 mM to induce expression of the *qrgB* allele. At 15, 30, and 60 minutes after IPTG was introduced, additional aliquots of culture were removed and processed as described above. For each timepoint, rifampicin treatment was allowed to proceed at 200 rpm shaking at 35°C for 10 minutes, at which time 40mL of rifampin-treated culture was added to each of two 50 mL conical tubes containing 1080 μL of fresh formaldehyde to induce crosslinking. Crosslinking was allowed to proceed for 5 minutes at room temperature with constant shaking, after which time the reaction was quenched by the addition of 8 mL of 2 M glycine. Quenched reactions were incubated for 10 minutes at room temperature with constant shaking, placed on ice for 10 minutes, then cells were collected by centrifugation for 5 minutes at 5,000 x g at 4°C. Crosslinked cell pellets from the matched 50 mL conical tubes were resuspended in 10 mL cold PBS and combined. The combined crosslinked cells were then harvested by centrifugation for 5 minutes at 5,000 x g at 4°C, resuspended in a second 10 mL PBS aliquot, and again harvested by centrifugation for 5 minutes at 5,000 x g at 4°C. The twice washed cross linked cell pellet was then resuspended in 400 μL of cold PBS, split to three 1.8 mL tubes and the final cell samples were collected via centrifugation for 5 minutes at 4°C, 10,000 x g. The supernatant was aspirated and pellets were flash frozen in an ethanol-dry ice bath, then stored at -80°C.

### RNA-seq

Total nucleic acid was extracted from cells, which had been lysed and preserved in DNA/RNA Shield and stored at -80°C, using an RNA Clean & Concentrator-96 kit (Zymo Research Corporation). Briefly, 300 μL of cell lysate was added to tubes containing 600 μL of RNA binding buffer and 900 μL of 100% ethanol, and vortexed. Each sample was bound to a filter (one well) of a 96 well spin column plate 700 μL at a time until all of the sample was bound. Washes were performed per manufacturer’s protocol. Samples were eluted using 25 μL of water.

DNA was removed by digestion with Baseline-ZERO DNase (LGC Biosearch Technologies). All of the samples (∼23 μL each) were added to DNA lo-bind tubes containing 60 μL of water and 10 μL of 10X Baseline-ZERO DNase Buffer. After vortexing, 2 μL of Murine RNase Inhibitor (NEB) and 5 μL of Baseline-ZERO DNase were added to each sample. After gentle mixing, tubes were incubated 37°C for 30 minutes.

Reactions were cleaned again using a Zymo RNA Clean and Concentrate Kit-96 (Zymo Research) according to the manufacturer’s instructions, and eluted with 25 μL of water. The QuantiFluor RNA System (Promega) was used to determine RNA concentration of samples.

For each sample, 0.5 µg of RNA was diluted to 11 µL and used as input material for NEBNext rRNA Depletion Kit for Bacteria (NEB). Depletion was performed per kit directions. Library preparation was performed using NEBNext Ultra II Directional RNA Library Prep Kit for Illumina (NEB) per kit directions for intact RNA (15 minute fragmentation) except that diluted (1:100) NEBNext Unique Dual Index UMI Adaptors DNA Set 1 were used instead of the NEBNext Adaptor for Illumina. Pooled libraries were sequenced using an Illumina NextSeq platform with 38 x 37 bp paired end reads.

### RNAP ChIP-seq

Crosslinked cell pellets were resuspended in 630 μL of lysis buffer [10 mM Tris pH 8.0, 1x cOmplete™, EDTA-free Protease Inhibitor Cocktail (Roche), 50 mM NaCl, 52 units/mL Ready-Lyse (Epicentre)] and incubated 15 minutes at 30°C. Samples were placed on ice, then sonicated in ice bath at 25% amplitude for 20 seconds (5 seconds on, 5 seconds off). To each sample, 5.4 μL of 100 mM MnCl_2_, 4.5 μL 100 mM CaCl_2_, 12 μL of RNase A (10mg/mL) and 12 μL of DNase I (Fisher) were added and mixed. Samples were incubated on ice for 30 minutes before 50 μL of 500 mM EDTA was mixed in to stop DNase I digestion. Samples were clarified by centrifugation (16k r.c.f.,10 minutes, 4°C), then liquid was split into new microfuge tubes (50 μL for input; 350 μL for RNA polymerase ChIP; ∼ 325 μL for other purposes). To each input sample, 450 μL of ChIP elution buffer (50 mM Tris pH 8.0, 10mM EDTA pH 8.0, 1% SDS) was added and mixed in. Inputs were maintained on ice until the end of the day (typically a few hours), then placed at 65°C for 12 hours to reverse cross-links.

After returning to room temperature, to each RNApol ChIP sample, an equal volume of 2X IP buffer (200 nM Tris pH 8.0, 600 mM NaCl, 4% Triton X100) with 2x cOmplete™, EDTA-free Protease Inhibitor Cocktail and 10 μL of purified anti-E. coli RNA Polymerase β Antibody (1μg/μL, BioLegend clone 8RB13) was added. ChIP samples were incubated overnight on a nutating platform at 4°C. After incubation overnight, 50 μL of Dynabeads protein G magnetic beads (Invitrogen; pre-washed once with 200 μL 1X IP buffer and resuspended in 1x IP buffer) were added to each ChIP sample and incubated 2 hours on a nutating platform at 4°C. Beads from ChIP samples were washed, in series, with 1 mL of Buffer A (100 mM Tris pH8, 250mM LiCl, 0.2% Triton x-100, 1mM EDTA), Buffer B (10 mM Tris pH8, 500mM LiCl, 0.1% Triton x-100, 1mM EDTA, sodium deoxycholate), Buffer C (10 mM Tris pH8, 500mM LiCl, 0.1% Triton x-100, 1mM EDTA), 1X IP buffer with 1 mM EDTA added, and 1X TE. Beads from ChIP samples were resuspended in 500 μL ChIP elution buffer, heated to 65°C for 30 minutes with 600 rpm shaking, and pulled out of solution by a magnetic stand. The eluted ChIP samples were transferred to new tubes and placed at 65°C for 12 hours to reverse cross-links.

After reversal of cross-links, all sample types were digested for 2 hours with RNase A (100μg/sample) and then 2 hours with proteinase K (200μg/sample). This was followed by phenol/chloroform extraction [add 500μL of phenol/chloroform/isoamyl alcohol (25:24:1), mix, separate layers (2 minutes at 21,130 r.c.f.), transfer water layer to new tube with 500 μL of chloroform/isoamyl alcohol (24:1), mix, separate layers (2 minutes at 21,130 r.c.f.), and then transfer water layer to new empty tube]. DNA from samples was precipitated by the addition of 1/25 volume of 5M NaCl, 2 μL of Glycoblue coprecipitant (Invitrogen), and 3 volumes of 100% ethanol, mixed, incubating 1 hour at 4°C, then >2 hr at -20°C. DNA was pelleted via centrifugation (16,000 r.c.f., 15 minutes, 4°C), washed once with ice-cold 95% ethanol, and air dried. Pellets were brought up in 0.1X TE (100 μL for input, 20 μL for IPOD or RNApol ChIP). QuantiFluor dsDNA dye (Promega) was used to determine DNA concentration prior to sequencing library preparation.

For each sample (input and ChIP), 0.5 ng of DNA was diluted to 50 µL and used as input material for NEBNext Ultra II DNA Library Prep Kit for Illumina (NEB) with NEBNext® Multiplex Oligos for Illumina Dual Index Primers Sets 1 and 2 (NEB). Manufacturer’s instructions were followed except the post-ligation bead clean-up, which was replaced with Oligo Clean & Concentrator kit (Zymo Research Corporation; performed per manufacturer’s protocol except DNA was eluted with 16.2 μL of 0.1X TE).

### NGS data analysis

#### Read pre-processing

Illumina sequencing adapters were trimmed from reads using cutadapt v4.0^52^ with the following command line arguments: –quality-base=33 -a AGATCGGAAGAGC -A AGATCGGAAGAGC -n 3 –match-read-wildcards. Trimmomatic v0.39^53^ was then used to remove low-quality bases from reads with the following command line arguments: - phred33 TRAILING:3 SLIDINGWINDOW:4:15 MINLEN:10.

#### RNAP ChIP-seq analysis

##### Read alignment, fragment counting, and enrichment inference

To align pre-processed reads, we first created a custom reference sequence comprising GenBank CP064350.1 (*V. cholerae* C6706 chromosome I), GenBank CP064351.1 (*V. cholerae* C6706 chromosome II), and the sequence of the plasmid pBRP02, which was used to overexpress QrgB. Alignment of pre-processed reads to the bowtie2 index created from this reference sequence was performed using bowtie2 v2.4.4^54^ with the following command line arguments: -q –end-to-end –very-sensitive –phred33 –fr -I 0 -X 2000 –dovetail. Position-sorted bam files were prepared using samtools v1.9^55^. Aligned fragments were counted at 5-bp resolution and the resulting bedgraph files were used as inputs to Enricherator to infer RNAP ChIP-seq enrichment^56^. Arguments of note to our call of “cmdstan_fit_enrichment_model.R” were --ext_subsample_dist 75 -- ext_fragment_length 150 --input_subsample_dist 70 --input_fragment_length 140 -- frac_genome_enriched 0.2 and --grad_samps 2.

We used Enricherator v0.3.0^56^ to infer RNAP occupancy and to estimate the contrast between occupancy in QrgB vs. QrgB*. Example code used for fragment counting and Enricherator analysis can be found at https://github.com/jwschroeder3/cdgc_code.

#### RNA-seq analysis

Our analysis of RNA-seq data is described in detail below, but briefly, we assembled the *V. cholerae* C6706 transcriptome using Rockhopper^57^, and quantified transcript abundance and differential expression in QrgB vs. QrgB* using a combination of kallisto ^58^ and the R package sleuth^59^.

##### Preparation of kallisto index

A kallisto index was prepared by first performing a reference-guided assembly of the transcriptome in our RNA-seq samples using Rockhopper [48]. Rockhopper was run using each chromosome, CP064350.1 (*V. cholerae* C6706 chromosome I) and CP064351.1 (*V. cholerae* C6706 chromosome II), as the reference separately, after which we pooled the two chromosomes’ Rockhopper assembled transcripts and operons. Some annotated transcripts present in the GenBank gff files for CP064350.1 and CP064351.1 were not assembled by Rockhopper, so we supplemented the transcript and operon assemblies from Rockhopper with any annotations in the GenBank gff files that were absent from the Rockhopper results, resulting in a final set of annotated and newly assembled transcripts and operons. In order to accurately quantify transcript abundance in antisense RNAs, for each CDS or operon of CDSs, we inserted a line into the final annotations representing a full-length antisense transcript for the given CDS or operon of CDSs. This procedure allowed kallisto to properly allocate antisense reads between short antisense RNAs and background fragments pseudoaligning to the full-length antisense annotations we inserted. Our manual inspection of alignments of RNA-seq data revealed the *snrC* locus to be on the minus strand of ChrII, from nucleotides 796,238 to 796,567, inclusive, and we inserted the *snrC* annotation into our annotations file accordingly. Finally, we extracted the sequence of each transcript using bedtools getfasta with the -s flag set to enforce stranded retrieval of each transcript’s sequence. The resulting fasta file was used to prepare a kallisto index for pseudoalignment.

##### RNA-seq quantification

Transcript abundance was estimated using kallisto v0.50.1 via the “kallisto quant” command with the --bootstrap_samples argument set to 100. The R package sleuth v0.30.1 was used to estimate differential expression of QrgB vs. QrgB*. Example code used for transcript assembly and differential expression analysis can be found at https://github.com/jwschroeder3/cdgc_code.

## Data availability

RNAP ChIP-seq data and enrichment scores are available under GEO accession GSE274215, and RNA-seq data and differential expression analyses are available under GEO accession GSE274216.

## Acknowledgements

We thank Jasper Gomez for the construction of C6706 *lac*-strain. We thank Sean Crosson and Hunter North for access and assistance in using Leica DMI 6000 microscope. We thank Lijun Chen and Anthony Schilmiller of MSU Mass Spectrometry core for their technical support. The research was supported by NIH grant R35 GM128637 (to L.F.), NSF grant AWD2025426 (to L.F.), and NIH grants GM139537, AI158433, and R21AI190733 and a MSU Tetrad award to C.M.W.

