## Supplementary material for "Negative feedback of cyclic di-GMP levels optimizes switching between sessile and motile lifestyles in *Vibrio cholerae*": Response to Reviewers

We thank the reviewers for their time and efforts on this manuscript. We apologize for the delay in submitting a revised manuscript, which was delayed by people leaving and health issues. We have addressed all the reviewers comments and included a point-by-point response below. Importantly we have added key new pieces of data that support and strengthen our conclusions:

1. **Fig. S1A-S1B-**We show decreased levels of c-di-GMP in WT cells (carrying the empty vector control) is similar to QrgB*. Thus, the decreased levels of c-di-GMP are due to the cells increasing in density instead of the presence of QrgB*, which addresses a key point made by two reviewers.
2. **Figs. 3B, 3C, and 3D-**In addition, we show that the *cdgC* expression dynamically changes in response to c-di-GMP levels using (i) promoter-luciferase assays (Fig. 3B) and (ii) analyzing *cdgC* expression using RT-qPCR (Figs. 3C-D).
3. **Fig. S3**-We demonstrated that *cdgC* expression was induced by another DGC, VC2224, suggesting this regulation occurs in response to increasing c-di-GMP concentrations and is not DGC specific.
4. **Fig. S4A-C-**We confirm the presence of the *cdgC* counter-transcribed sRNA, *snrC*, in the Δ*vpsR* mutant by strand specific transcript quantification using RT-qPCR (Fig. S4A). We further show by RT-qPCR that *snrC* is significantly upregulated in Δ*vpsR* mutant compared to the WT (Figs. S4B-C).

We hope these changes have made the manuscript ready for publication.

A full point-by-point response to reviewer comments is included below:

**Reviewer #1 (Evidence, reproducibility and clarity (Required)):**

The study by Rangarajan, Schroeder et al. uncovers a negative feedback loop that controls cdG levels in Vibrio cholerae. They demonstrate that this feedback loop requires VpsR, one of the master regulators of cdG-dependent responses in this organism. Furthermore, they show that mutating the phosphodiesterase CdgC eliminates this negative feedback loop. This research is timely and addresses an important question about the regulation of this important secondary messenger molecule in bacterial species. The manuscript is well written and clearly presented. However, some of the results are fairly preliminary and there are alternative models that could explain the results obtained. Also, I disagree with the authors interpretations of some of their data. Specific points for the authors to consider are discussed below.

**RESPONSE 1:** We thank the reviewer for their careful reading of the work, their overall supportive assessment of the impact of the work, and their many useful suggestions. We appreciate the fact that the reviewer finds the work to be “timely” and “important”. We will address each of their major points below.

Line 113-115: This reduction in cdG levels observed with expression of QrgB* is quite striking. In fact, the ratiometric "increase" in cdG levels observed for QrgB/QrgB* is largely driven by the decrease in cdG levels in the QrgB* condition as opposed to an increase in cdG in QrgB. Are the authors suggesting that the decrease observed with QrgB* is due to the natural reduction of cdG levels due to quorum sensing and the transition of cells to the high cell density state? What is the change in OD600 during this time course? Alternatively, isn't it possible that the expression of QrgB* is causing a response that results in a decrease in cdG? If so, this would change a lot of the conclusions in the present study. Perhaps one way to address this is if the authors have data on the cdG levels in wildtype cells during a similar time course without any induction of QrgB or QrgB*.

**RESPONSE 2:** We appreciate the reviewer's concern about the decreased cyclic di-GMP in QrgB* mutant. However, it is known from the previous studies that cells at stationary phase possess decreased cyclic di-GMP levels when compared to the exponential phase cells (PMID: 24799624, Figure 2). It is also shown that cyclic di-GMP level is higher in the locked low cell density mutant when compared to the wild-type upon overexpression of empty vector, QrgB and QrgB* indicating high cell density effect of decreased cyclic di-GMP levels (PMID 18223081, Figure 4B). We have added “To address whether expression of QrgB* results in decreased levels of c-di-GMP, we transformed an empty plasmid in the wild-type strain and checked the levels of c-di-GMP levels. We observed the c--di-GMP levels decreased in wild-type cells with empty vector as the OD increased after the addition of IPTG, similar to QrgB* **(new Fig. S1A).”** This indicates that expression of QrgB* does not decrease the levels of cdG levels, but this decrease is due to the cells transitioning into the high cell density quorum state. This is consistent with the known reduction of c-di-GMP levels when cells are in high cell density^22,24^”.

To further validate the quantification of c-di-GMP concentrations that we performed, we quantified the area of cells in different c-di-GMP states to ensure that the differences we were observing could not be accounted by this variable. No significant difference in the area of cells in any condition analyzed was observed (new **Fig. S1B**).

Fig 2B: The authors indicate that the results with the PcdgC reporter confirm their RNA-seq results. I don't agree with this based on my interpretation of the presented data. While statistical comparisons are not provided, the relative change in PcdgC reporter activity when comparing 0 min to 60 min is very modest for all samples. This is especially relevant for the WT+QrgB condition, which should see a marked increase in reporter activity based on the RNA-seq results presented in 2A. The potential post-transcriptional regulation of CdgC discussed in the subsequent figures further complicates this. The data in 2C suggest that cdgC is associated with a negative feedback circuit that keeps cdG levels low in the wildtype as shown in 1A. But the mechanism underlying how this actually occurs remains unclear. Defined reporters or direct analyses of cdgC transcription and translation would be valuable to elucidate this.

**RESPONSE 3:** We appreciate the reviewer’s point here and agree that additional quantifications of *cdgC* expression with a higher dynamic range are warranted. In our revised manuscript, we generated a *cdgC*-luciferase transcriptional reporter that exhibits robust induction by c-di-GMP in the wild type background by not the Δ*vpsR* mutant (**new Fig. 3B**). This reporter only measures transcription initiation, thus clearly demonstrating that VpsR is acting to induce transcription of *cdgC* in response to elevated c-di-GMP. We further confirmed the RNA-Seq and *cdgC-lux* fusion via RT- qPCR in both the wild type and Δ*vpsR* mutant (**new Fig. 3C and 3D**). Both approaches are consistent with our initial RNA-Seq observations.

The data in 2C show that mutating cdgC now allows for similarly high levels of cdG in parent vs ∆vpsR. But how specific is this response to cdgC? An alternative possibility is that elevating the baseline cdG level is sufficient to bypass this feedback loop. For example, CdgJ is a phosphodiesterase that has previously been characterized in Vibrio cholerae and cdgJ mutants exhibit phenotypes consistent with elevated cdG. Performing similar experiments in the cdgJ mutant to show that the results seen in 2C do not occur when another phosphodiesterase is mutated would strengthen the conclusion that CdgC is specifically required for the negative feedback circuit that controls cdG levels.

**RESPONSE 4:** We agree with the reviewer’s point that other manipulations of *V. cholerae* can alter c-di-GMP concentrations. However, our argument is that under the conditions of induced increases in c-di-GMP levels in our experiments, CdgC is the key PDE that mediates the negative feedback response. This is evidenced by the fact that *cdgC* is the only PDE or DGC specifically regulated by VpsR, the negative feedback response is lost in a Δ*cdgC* mutant, and restoration of *cdgC* restores this response.

To experimentally address whether *cdgC* induction was a general response to increased c-di-GMP concentrations (versus QrgB induction specifically), we overexpressed a strong DGC of *V. cholerae,* VC2224, and found via RT-qPCR that *cdgC* was induced 4-fold in the wild-type but not the Δ*vpsR* mutant (**new** **Fig. S3**), indicating that *cdgC* responds to general increase in the c-di-GMP levels irrespective of the the DGC. This conclusion is supported by several published transcriptomic studies in different conditions that demonstrate *cdgC* expression is upregulated during increase c-di-GMP (PMID: 17071756, 25622616, 35302986).

The only data that link CdgC to the negative feedback circuit are the assays performed in 2C that directly measure intracellular cdG levels. But the authors make a compelling case for the negative feedback circuit in generating a pulsatile response to QgrB induction in the wildtype in 1F. if CdgC is required for this negative feedback response, then a CdgC mutant should lose this pulsatile response. This should be tested to further support the model presented.

**RESPONSE 5:** We agree with the reviewer and indeed directly demonstrate this in Fig. 2E (blue solid line), showing that VpsR does not substantively impact c-di-GMP dynamics in a *cdgC* mutant or a *cdgC* overexpression strain.

Fig. 3B: It would be valuable to also plot the sense transcripts in this same region so that the expression of cdgC can be directly compared to the expression of SnrC. Also, can the authors speculate on why the RNAP enrichment at PsnrC in 3A does not reflect the expression levels of SnrC observed in 3B?

**RESPONSE 6:** We thank the reviewer for making these suggestions. We have replotted the corresponding data (which appears in Fig. S4 of the revised manuscript) to show both the plus and minus strand RNA abundances in this region. We also comment in the text on the RNAP occupancy, which does in fact show a small peak at the likely SnrC promoter at the last *vpsR* timepoint:

A concomitant increase in RNA polymerase occupancy near the likely snrC transcription start site can be seen at the 60-minute timepoint in the Δ*vpsR* strain (Fig. 3A; position 0.7965 Mbp), although the occupancy is relatively weak likely due to the fairly low transcript levels seen for SnrC under this condition.

Line 173-174: I am not sure how the authors are concluding that VpsR induces SnrC. I believe their data in 3B indicates the exact opposite, which is that SnrC expression is negatively regulated by VpsR. This finding is also not reflected in Fig.5.

**RESPONSE 7:** We thank the reviewer for identifying this typo and agree with their conclusion that VpsR represses *snrC* expression.

.

The data presented in Fig. 3 are compelling, but they only suggest that SnrC may be an anti-sense small RNA. More rigorous analyses are needed to make any firm conclusions about whether this is a bona fide sRNA and to define its potential mechanism of action. Northern blots should be performed to directly assess SnrC levels. Also, can the PsnrC promoter be mutated to prevent production of this small RNA while leaving cdgC function intact? This would allow for more rigorous genetic analysis to assess the impact of SnrC on CdgC transcription and translation. At the very least, claims of SnrC as a small RNA need to be toned down

**RESPONSE 8:** To address whether *snrC* was present, we performed a strand specific q-RT-PCR to detect SnrC. Our results showed that the wild type strain had quantifiable *snrC* signals whereas a Δ*cdgC* mutant lost all detectable *snrC,* confirming the presence of this RNA in the cell. Moreover, we quantified higher levels of *snrC* in the Δ*vpsR* mutant compared to the wild type, but regulation by c-di-GMP was not observed (**new Fig. S4D**). Given that we can quantify *snrC* and demonstrate that it modulates c-di-GMP in a *cdgC*-dependent manner, we argue that it fits all the criteria necessary to define this as a sRNA. However, the regulation of *snrC* appears to be complex, and given that it is not central to the negative regulation that is the focus of this manuscript, we feel any other experiments are out of scope for this work but would certainly be interesting in follow-up studies. To this end, we have reduced discussion of *snrC* and moved the data related to it to a supplemental figure. We have also removed *snrC* from the final model proposed in Fig. 5.

Fig. 4C: this model system and data are very interesting. But they do not necessarily indicate that it is the negative feedback circuit that CdgC participates in that is important for these transitions. They may simply reflect that elevated cdG results in a defect in these assays. As mentioned above, cdgJ mutants have previously been shown to exhibit phenotypes consistent with elevated cdG. So, what is the phenotype of a cdgJ mutant in these assays? Since the cdgJ mutant should still have CdgC-dependent negative feedback control of cdG levels, this mutant may not have any defect. This result would be particularly compelling and support the authors conclusions that CdgC-dependent feedback control of cdG is critical for balancing biofilm formation and dispersion.

**RESPONSE 9:** Indeed, we agree with the reviewer and predict that deletion or overexpression of another enzyme involved in equilibrium of c-di-GMP would also impact the dynamics of the system studied here. The point of our experiments, rather, is that CdgC plays an important role in the context of the intact, wild type regulatory network due to its unique responsiveness to c-di-GMP levels (Fig. 2A), and we believe that our data are certainly sufficient to demonstrate that CdgC has an important impact on the phenotypes discussed in the manuscript. Given that our strain of *V. cholerae* has 62 enzymes predicted to impact c-di-GMP, we would expect many of these, including CdgJ, would have a fitness defect in this assay.

Line 299-302: This is an interesting hypothesis, and could be tested with a cdgC catalytic site mutant. Or by otherwise inactivating CdgC phosphodiesterase activity while leaving the cdgC gene largely intact so that PsnrC expression can still be monitored. Again, these types of experiments are needed to make any firm conclusions about SnrC.

**RESPONSE 10:** We thank the reviewer for this interesting experiment, but as stated in Response 8 we have limited our discussion of *snrC* and this hypothesis is no longer proposed in the revised manuscript.

**Reviewer #2 (Evidence, reproducibility and clarity (Required)):**

The work from Rangarajan, Schroeder et al. examines dynamics of c-di-GMP (cdG) gene regulation and homeostasis in Vibrio cholerae following cdG overexpression. Relative increase in cdG levels are stimulated by the heterologous QrgB DGC enzyme, and the resulting changes in gene expression (biofilm up, motility down) are largely due to the cdG-dependent transcriptional activator. VpsR induction leads to activation of GGDEF/EAL protein CdgC; only its EAL (cdG-degrading) domain is active, and this activation leads to a feedback loop that results in a reduction in cdG levels. ChIP-Seq with RNAP led the authors to an unannotated sRNA that is transcribed within the cdgC ORF in the antisense direction that they name sncR, and whose expression is positively regulated by cdG. Expression of sncR negatively regulates cdG levels through CdgG.

The authors could thus break the negative feedback loop they discovered and generate sustained high levels of cdG in the cell using ΔvpsR or ΔcdgC single or double mutants, and in those cases they observe growth defects.

Overall, the experiments presented are high quality and the results offer significant novelty. The RNA-seq studies in Fig. 1 provide higher resolution than previous studies, including following the early dynamics upon cdG induction and resolving the VpsR/VpsT-dependence of individual genes and classes of genes. The figures are exceptionally clear, and the text is accurate and well cited/referenced. Multiple instances of the text do require additional clarification as noted below.

**RESPONSE 11:** We appreciate the reviewer’s overall positive assessment, supportive comments, and careful reading of the manuscript. We have addressed the suggestions raised by the reviewer as detailed below.

Major comments:

1. High levels of sncR are identified only in a ΔvpsR strain (Fig. 3B), yet the authors' model is that VpsR is not directly regulating sncR expression (Fig. 5). Instead, they suggest that the VpsR-dependence of the phenotype is due to only observing the highest levels of cdG in the absence of VpsR. If there is a way to clarify the connection of VpsR and sncR that would strengthen the manuscript.

**RESPONSE 12:** Similar concerns were mentioned by Reviewer #1 which we addressed in RESPONSE 8. To summarize, we used Q-RT-PCR to demonstrate that *snrC* can be detected in *V. cholerae* and we found its expression was higher in the absence of VpsR, but the role of c-di-GMP in this response remains enigmatic. Thus, we place less emphasis on *snrC* in the revised manuscript.

2. In Fig. 4BC, are these strains similarly vpsL- as most of the paper? If so, is it expected that they can form biofilms? I would recommend explaining this assay further, or if they are vpsL+ then clarifying in the text.

**RESPONSE 13:** We thank the reviewer for the suggestion. As noted in the Methods section of the original manuscript, the strains used for the biofilm assay in Figure 4C are indeed *vpsL^+^*. We have added additional explanatory text to the text accompanying Fig. 4C to clarify this point.

3. L106-116: In Fig. 1A, why does QrgB overexpression not lead to elevated cdG at 15 m except upon normalization to QrgB* (Fig. 1C)? Is this connected to the feedback loop characterized in the manuscript (does it act that quickly?) or due to another effect?

**RESPONSE 14:** As we note in RESPONSE 2 to Reviewer #1, the basal levels of c-di-GMP in *V. cholerae* are decreasing during this period of the growth curve, which we further demonstrate in the new data quantifying c-di-GMP in the vector control strain presented in Fig. S1A. Thus, although QrgB induction does not appear to increase c-di-GMP in Fig. 1B in the wild type strain (i.e. a relatively flat line), in essence it is significantly increasing c-di-GMP as the overall levels are decreasing.

Related, the authors attribute the decrease in the denominator (QrgB* levels; Fig. 1B) to the cells transitioning to the high cell density state. Is it clear that this would occur in the time frame of 60 min?

**RESPONSE 15:** Yes, we see a similar decrease in the vector control strain as described in RESPONSE 2 and RESPONSE 14.

Minor comments:

4a. In Fig. 1D, what does the null symbol represent?

**RESPONSE 16:** In the revised manuscript, we now clarify that the null symbol in Fig. 1D indicates that data for a given gene are unavailable due to deletion of the gene in question in that experiment.

4b. L200-203, more explanation would benefit the assertion of the Type II error proposed. Are the authors claiming this is noted as not significant due to the high number of multiple comparisons?

**RESPONSE 17:**

We had been alluding to the possibility of false negatives, which are fairly common in RNA-seq data, but in any case this text no longer appears in the revised narrative**.**

**Reviewer #3 (Evidence, reproducibility and clarity (Required)):**

This study investigated c-di-GMP (cdG)-dependent gene expression in Vibrio cholerae, an area that has been thoroughly researched in earlier studies. The transcriptomics results of the first sections largely confirm what has been previously described, but identify an unexpected, sudden c-di-GMP drop at OD600=1, which may be either the result of a feedback loop, as the authors suggest, or an experimental artifact (as explained in Major concerns). The most intriguing result of the study, in my opinion, is the identification of a potential antisense RNA, designated SnrC, that may counteract expression of the main cdG phosphodiesterase gene, cdgC. Unfortunately, the data are insufficient to conclude that SnrC is real. The follow-up competitive analysis of the cdgC mutants and growth analysis of the mutants with super-high cdG levels are not insightful because of low biological relevance. Overall, while some intriguing leads are described in this work, major experimental concerns exist and some conclusions may have to be revised.

**RESPONSE 18:** We must respectfully disagree with the reviewer’s overall assessment that the work is not impactful and would indeed point to the comments of Reviewer’s 1 and 2 that state otherwise. We address the reviewer’s other comments in the specific point by point responses below.

MAJOR CONCERNS

1. The experimental setup may have had a major flaw. A sudden drastic drop in c-di-GMP levels at OD600=1 may be due to sampling/inducer addition. Until this is solved, it impossible to evaluate the adequacy of conclusions based on transcriptomics analysis.

Fig. 1B shows an approx. 3-fold drop in c-di-GMP levels within 15 min upon induction of the QrgB-mutant protein expression. This drop cannot be explained by "transitioning to the high-cell density quorum state", as suggested by the authors, because such a transition likely takes more than 15 min, and it does not start so abruptly at precisely OD600=1 (when IPTG was added). A different, more benign explanation is more likely. Below are a couple of ideas to investigate.

- Can the cdG decrease be due to the sudden drop in dissolved oxygen levels during the sample withdrawal for c-di-GMP measurements and IPTG addition? If so, the cultures were experiencing first hypoxia, then (upon restoration of shaking) oxidative stress, which affect the c-di-GMP transcriptome. I suspect that an approx. 3-fold drop in cdG levels does not take place in the undisturbed cultures that are allowed to grow for 15 min after reaching OD600=1.

**RESPONSE 19:** Given that all three reviewers commented on these data, we think this is an important point to address, and have detailed this in RESPONSE 2 (see above). The bottom line is that the decrease of c-di-GMP during these time points is not specific to QrgB* but the natural response of *V. cholerae.* Whether this is due to quorum sensing or some other mechanism as suggested by the reviewer is irrelevant to the conclusions of our manuscript. We do not think that the c-di-GMP begins to drop at an OD_600_ of 1.0 as suggested by reviewer, but rather it is continuously dropping as the cells transition to higher density as we are just viewing a few discrete snapshots rather than the entire curve.

- Does the mutant QrgB* protein contain a cdG-binding site? If so, QrgB* may soak up intracellular cdG, whereas the intact QrgB may counteract the soaking effect by synthesizing cdG.

**RESPONSE 20:** Yes, QrgB does have a c-di-GMP binding I-site, but as we now show in Fig. S1A the c-di-GMP levels when expressing this protein match a vector control, suggesting this binding site is not significantly impacting c-di-GMP concentrations in our experiment conditions. Moreover, we have used the QrgB* active site mutant in many experiments and new observed a significant biological effect that suggests binding of c-di-GMP to this protein impacts c-di-GMP signaling. Two published examples are shown below.

In the paper by Pursley, Maiden, and Waters (PMID: 29311281), the data in Fig. 4A measures induction of four promoters in response to increase c-di-GMP levels by inducing QrgB or QrgB*. As you can see, there are clear effects upon addition of the inducer, IPTG, with QrgB whereas no effects are observed upon IPTG induction of QrgB*, suggesting that QrgB* has no effect on available c-di-GMP levels or c-di-GMP signaling in *V. cholerae.*

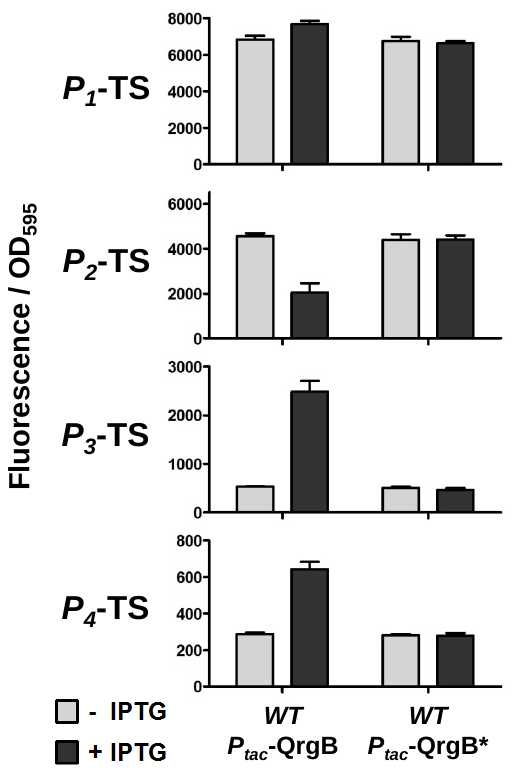

Similarly, in Fernandez et. al. (PMID: 29311281), in Fig. 2 increasing IPTG induction of QrgB leads to a reduction in cell curvature whereas increasing IPTG induction of QrgB* has no effect.

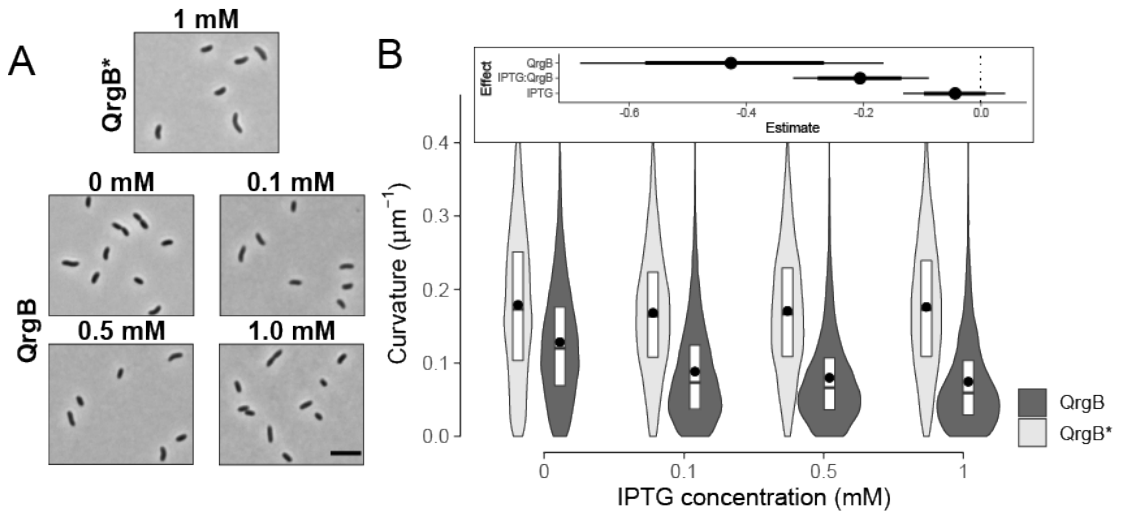

Both of these pieces of evidence (and many more experiments that we have done) demonstrate that QrgB* on its own does not impact c-di-GMP signaling.

2. Evidence of the potential existence of the antisense RNA, SnrC, is weak.

It is not convincing that snrC is real despite the notion that bioinformatics data lack statistical significance (l. 197-203). A functional test, where an overexpressed antisense RNA lowers expression of the sense RNA (Fig. 3D) is obviously insufficient because this would work for any gene, whether antisense RNA is real or not. To verify that SnrC is real, one needs to show, at least, a snrC RNA band on a Northern blot and/or results of the snrC transcriptional fusion.

**RESPONSE 21:** We have addressed the concerns regarding *snrC* in RESPONSE 10 and now show that *snrC* exists using RT-qPCR. We think it is debatable whether expression of any countertranscribed RNA would reduce expression, but this is beside the point as we demonstrate that *snrC* IS produced and it DOES impact c-di-GMP concentrations in a *cdgC* dependent manner. We completely agree with the reviewers that our understanding of *snrC* is at the early stages and have thus moved all data regarding it to a supplemental figure and removed it from the final model as it is not relevant for the VpsR/CdgC negative feedback loop presented here.

ADDITIONAL CONCERNS

l. 95: More information about QrgB is needed, eg, is QrgB a cytoplasmic or membrane protein; does its DGC activity depend on interactions with small molecules or other proteins; does V. cholerae contain a QrgB homolog; does it have a cdG-binding site?

**RESPONSE 22**: We have added more information about QrgB as described below. Importantly, QrgB has been used to modulate c-di-GMP levels in *V. cholerae* and many other bacterial species, and we have demonstrated that it increased c-di-GMP concentrations to physiological levels similar to the low-cell density state (PMID: 29311281).

“QrgB is a cytoplasmic protein containing N-terminal sensor PAS domain and C-terminal GGEEF domain. It also contains RxxD I-site in the GGEEF domain that binds cyclic di-GMP. QrgB does not have any homology to *V. cholerae* proteins which negates the possibility of interaction with *V. cholerae* proteins”.

l. 106: It is erroneous to suggest that in the wild type, c-di-GMP levels increased at 15 & 30 min when they were largely unchanged (Fig. 1A).

**RESPONSE 23:** We have addressed this point in RESPONSE 2 and 14, but the bottom line is QrgB increased c-di-GMP levels relative to the QrgB* (or vector) control strains in which c-di-GMP is decreasing. Thus, we would argue this statement is not erroneous, and we have made substantive clarifications to justify it in the revised manuscript.

l.143-52: That cdgC may be part of the negative VspR-mediated feedback on cdG levels is evident from refs. 11,13,14 and from the confirmation data in Fig. S1. I'm afraid it is not at all clear from Fig. 2A that cdgC is "the sole clear candidate for negative feedback on cdG levels". Please provide more substantive reasoning.

**RESPONSE 24:** Our emphasis on cdgC arises because in our RNA-seq experiments, of all the proteins encoding the pertinent domains, CdgC was the only one to show consistent, increasing levels over the course of QrgB induction in both the WT and *vpsT* backgrounds (data in Figure 2A). We further demonstrate that the VpsR negative feedback effect is lost in a Δ*cdgC* mutant or when *cdgC* is overexpressed, demonstrating that in our conditions *cdgC* is the sole factor mediating the observed negative feedback effect. We certainly do not claim to be the first to demonstrate VpsR regulation of *cdgC* and indeed cite all the relevant references. But we feel this study contributes to prior knowledge by a more kinetic analysis of c-di-GMP levels in the wild type and Δ*vpsR* mutant strains, and, most importantly, demonstrating that disruption of this negative feedback response has a fitness cost both in terms of c-di-GMP toxicity and flexibility transitioning from sessile to motile states.
